# Role for Merlin/NF2 in transcription elongation through interaction with the PAF complex

**DOI:** 10.1101/717769

**Authors:** Anne E. Roehrig, Kristina Klupsch, Juan A. Oses-Prieto, Selim Chaib, Stephen Henderson, Warren Emmett, Lucy C. Young, Silvia Surinova, Andreas Blees, Anett Pfeiffer, Maha Tijani, Fabian Brunk, Nicole Hartig, Marta Muñoz-Alegre, Alexander Hergovich, Barbara H. Jennings, Alma L. Burlingame, Pablo Rodriguez-Viciana

## Abstract

The PAF complex (PAFC) coordinates transcription elongation and mRNA processing events and its CDC73/parafibromin subunit functions as a tumour suppressor. The NF2/Merlin tumour suppressor functions at the cell cortex and nucleus and is a key mediator of contact inhibition. Here we provide a direct link between nuclear Merlin and transcription elongation controlled by cell-cell adhesion. Merlin interacts with the PAFC in a cell density-dependent manner and tumour-derived inactivating mutations in both Merlin and CDC73 mutually disrupt their interaction. Growth suppression by Merlin requires CDC73 and Merlin regulates PAFC association with chromatin in a subset of genes. We also identify by CDC73 affinity-proteomics a role for FAT cadherins in regulating the Merlin-PAFC interaction. Our results suggest that in addition to its function within the Hippo pathway, nuclear Merlin is part of a tumour suppressor network which coordinates postinitiation steps of the transcription cycle of genes mediating in contact inhibition.

## INTRODUCTION

Normal cells cease to proliferate when they come into contact with each other and assemble intercellular junctions. Contact-dependent inhibition of proliferation is essential for normal tissue homeostasis and its loss is one of the hallmarks of cancer (Hanahan and Weinberg, 2011; McClatchey and Yap, 2012). The molecular mechanisms underlying contact inhibition remain unclear but Merlin, the protein encoded by the *NF2* tumour suppressor gene is known to be an important mediator (Cooper and Giancotti, 2014).

Germline mutations in *NF2* cause Neurofibromatosis type 2, a cancer syndrome characterized by the development of schwannomas, meningiomas and ependymomas. Somatic *NF2* mutations are frequently found in these tumour types as well as at lower frequency in several others, suggesting that *NF2* has a broad tumour suppression function (Cooper and Giancotti, 2014; Dalgliesh et al., 2010; Forbes et al., 2015).

Merlin functions as an upstream regulator of the Hippo tumour suppressor pathway to inactivate the YAP/TAZ transcriptional co-activators (Li et al., 2014; Yin et al., 2013; Zhang et al., 2010). In the absence of Hippo signalling, YAP/TAZ translocate to the nucleus to promote transcription of genes that regulate tissue homeostasis, organ size and tumorigenesis (Ma et al., 2019). Merlin has also been implicated in controlling the surface availability of membrane receptors and in the regulation of multiple signalling pathways (Li et al., 2012).

Merlin can localize to different cellular compartments. It is found at the cell cortex in lamellipodia and membrane ruffles and at cell–cell contacts in dense cultures where it can interact with α-catenin and Par3 at nascent adherens junctions and with AMOT/Angiomotin at tight junctions (Gladden et al., 2010; Yi et al., 2011). Merlin can also exhibit a punctate distribution that has been attributed to intracellular vesicles or particles that can move along microtubules in a kinesin and dynein-dependent manner (Bensenor et al., 2010; McCartney and Fehon, 1996; McClatchey and Giovannini, 2005). In addition, Merlin can also localize to the nucleus in a density and cell cycle dependent manner (Furukawa et al., 2017; Gronholm et al., 2006; Kressel and Schmucker, 2002; Li et al., 2010; Muranen et al., 2005).

Transcription of mRNAs is a tightly regulated process consisting of discrete stages of RNA polymerase II (Pol II) recruitment, initiation, elongation and termination. Transcription elongation is now recognized to play a key role in regulated gene expression. In particular, promoter-proximal pausing of Pol II and its release to initiate productive elongation are key regulatory steps in transcription with its precise intensity and the mechanisms regulating pause release likely varying from gene to gene (Adelman and Lis, 2012; Jonkers and Lis, 2015; Nechaev and Adelman, 2011). After promoter-proximal pause release, elongation rates also vary between genes and are closely linked to co-transcriptional RNA processing steps such as splicing, cleavage and polyadenylation (Jonkers and Lis, 2015). Chromatin represents a barrier to transcription by Pol II that has to be overcome as well as restored following polymerase passage. Accordingly, elongation is also tightly coupled with modulation of chromatin structure through the concerted actions of chromatin remodelers, histone-modifying enzymes and histone chaperones (Petesch and Lis, 2012).

The Pol II-associated factor 1 complex (PAFC) has been linked to multiple transcription related processes including communicating with transcriptional activators, recruiting histone modification factors, release from promoter-proximal pausing, facilitating elongation across chromatin templates, recruitment of 3’ end processing factors and mRNA stability (Chen et al., 2015; Jaehning, 2010; Jo et al., 2014; Yang et al., 2016; Yu et al., 2015). In humans it is composed of 5 subunits, PAF1, CTR9, LEO1, CDC73 and SKI8. CDC73, also called Parafibromin, is inactivated in Hyperparathyroidism-jaw tumour (HPT-JT) syndrome, an autosomal dominant disorder characterized by the development of parathyroid tumours as well as uterine and renal cancers. (Newey et al., 2010). The PAF1 subunit localizes to 19q13, a locus amplified in several tumours and has transforming properties highlighting a role for the PAFC in cancer (Chaudhary et al., 2007).

Here we show that Merlin interacts with multiple proteins involved in transcription elongation and mRNA processing including the PAFC, the CHD1 chromatin remodeller and the TAT-SF1 elongation/splicing factor suggesting a role of Merlin in coordinating transcription elongation and mRNA processing. We use genomics to identify Merlin-regulated genes and proteomics to show Merlin can regulate the PAFC interactome and identify an interaction between CDC73 and FAT cadherins that is required for Merlin interaction with the PAFC. Our results suggest that Merlin is part of a tumour suppressor network which helps regulate expression of genes mediating contact inhibition by coordinating initiation and post-initiation steps of their transcription cycle in response to cell adhesion.

## RESULTS

### Merlin interacts with proteins that function in transcription elongation and splicing

To identify Merlin interacting proteins that may shed light on its function, overexpressed Merlin was affinity purified from HEK293T cells and co-purifying proteins identified by mass spectrometry (Figure 1A and S1A). We identified several known Merlin-binding partners such as the VPRBP/DCAF1 and DDB1, members of a Cullin 4 containing E3 ubiquitin ligase complex (CRL4^VPRBP^) (Huang and Chen, 2008; Li et al., 2010; Yi et al., 2011)and AMOTL1, a tight-junction associated protein (Huang and Chen, 2008; Li et al., 2010; Yi et al., 2011). We also identified here new proteins that co-purified specifically with Merlin (but not other control baits) such as: CDC73, PAF1, CTR9, LEO1, and SKI8, all core components of the PAFC; CHD1, a chromatin remodeller implicated in elongation (Simic et al., 2003); components of the U2 small ribonucleoproteins complex (U2 snRNP) of the spliceosome; TAT-SF1, an elongation factor linked to splicing (Fong and Zhou, 2001) and PELP1, a protein that interacts with several transcriptional activators and chromatin-modifying proteins and has also been linked to splicing (Chakravarty et al., 2010; Mann et al., 2014) (Figure 1A and S1A).

**Figure 1.**
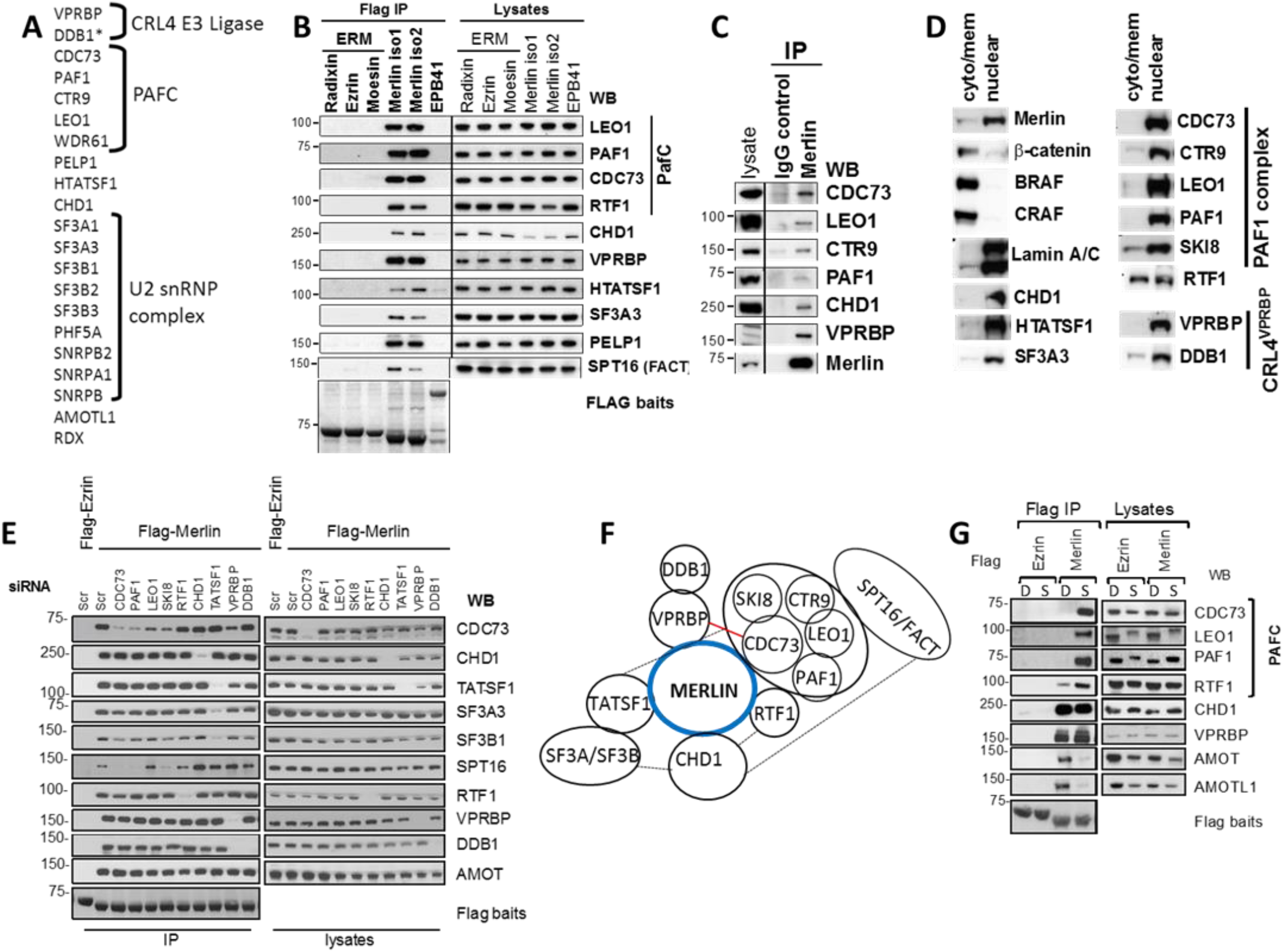
Merlin interacts with nuclear proteins that function in elongation and splicing. **(A)** Proteins identified by mass spectrometry after Merlin affinity purification from HEK293T cells. (see also Figure S1A). **(B)** Merlin isoforms 1 and 2 but not ERM proteins interact with proteins involved in transcription. Flag-tagged proteins were transfected in HEK293T cells and endogenous proteins detected on Flag IPs and lysates by western blot (wb). **(C)** Endogenous Merlin co-IPs with the PAFC, CHD1 and VPRBP. IPs of Merlin or control antibody from IOMM-Lee meningioma cells. **(D)** Merlin and its interactors reside in the nucleus. Western blot cytoplasmic/membrane and nuclear fractions from IOMM-Lee cells (see also Figure S1B). **(E)** Merlin independently binds to the PAFC, CHD1 and TAT-SF1. Flag-Ezrin or Merlin were purified from HEK293T cells transfected with the indicated siRNA oligos and interacting endogenous proteins detected by wb. **(F)** Schematic summarizing results from D. Dotted lines indicate interactions reported in the literature. Red line indicates interaction shown in Fig 4. Note that interactions between subunits of the PAFC are ‘arbitrary’. **(G)** Cell density inversely regulates Merlin’s interaction with the PAFC and AMOT proteins. Flag-Ezrin or Merlin were purified from either dense or sparse HEK293T cells. WB: Western blot, IP: Immunoprecipitation, Scr: scrambled control oligo, D: dense cells, S: sparse cells. See also Figure S1.

To validate the specificity of these interactions, we expressed Flag-tagged versions of Merlin, the three closely related ERM proteins, Ezrin, Radixin and Moesin, and the FERM-domain containing erythrocyte membrane protein band 4.1 (EPB41) in HEK293T cells. Immunoblotting of Flag-immunoprecipitates (IPs) showed that endogenous LEO1, PAF1, CDC73, CHD1, PELP, TAT-SF1 and SF3A3 interact specifically with both Merlin isoforms 1 and 2 but not with the three ERM proteins or EPB41 (Figure 1B). RTF1, which has been shown to physically and functionally associate with the PAFC (Kim et al., 2010), was not identified by mass spectrometry but could be detected by immunoblotting in Flag-IPs. Similarly, the SPT16 subunit of the histone chaperone FACT associated with Merlin. Importantly, endogenous Merlin co-immnuprecipitated with the PAFC components as well as CHD1 and VPRBP (Figure 1C), confirming that these interactions take place at physiological levels of Merlin.

Most of the novel Merlin interactors we identified are involved in transcription and this is consistent with the nuclear localization of a pool of Merlin (Furukawa et al., 2017; Kressel and Schmucker, 2002; Li et al., 2012; Muranen et al., 2005). In cell fractionation experiments endogenous Merlin can be readily detected in the nuclear fraction together with all the identified interactors (Figure 1D and S1D). Taken together these results suggest a novel role for nuclear Merlin in transcription through association with proteins associated with transcription elongation, splicing and chromatin remodeling.

In an effort to shed light on the nature of these Merlin interactions (e.g. direct vs indirect), we measured Merlin associations in Flag-Merlin IPs after knock-down with siRNAs of different interactors. Knockdown of TAT-SF1 or CHD1 had no effect on Merlin’s interaction with PAFC subunits (only CDC73 shown) and conversely, knockdown of PAFC subunits had no effect on Merlin’s interaction with TAT-SF1 or CHD1, suggesting the PAFC, TAT-SF1 and CHD1 bind independently to Merlin (Figure 1D). Knockdown of TAT-SF1 did inhibit the interaction with U2 snRNP complex components SF3A3 and SF3B1, consistent with Merlin binding the U2 snRNP complex indirectly via TAT-SF1, ((Fong and Zhou, 2001)). Knockdown of CDC73, PAF1 and SKI8 subunits of the PAFC, but not CHD1, inhibits the Merlin-SPT16 interaction, suggesting the FACT complex associating indirectly with Merlin through the PAFC (Figure 1E). In contrast, knockdown of PAFC subunits did not have any effect on binding of RTF1 and similarly knockdown of RTF1 had no effect on the interaction with PAFC subunits, consistent with then having independent roles in elongation (Cao et al., 2015; Mbogning et al., 2013). Knockdown of VPRBP only inhibited the interaction of Merlin with DDB1, consistent with DDB1 binding indirectly through VPRBP (Huang and Chen, 2008; Li et al., 2010). A model summarizing the data derived from these experiments is provided in Figure 1F.

Because of Merlin’s role in contact inhibition, the effects of cell density on Merlin’s interactions were studied. HEK293T, IOMM-Lee meningioma (Lee, 1990) and HMLE nontransformed mammary epithelial cells (Elenbaas et al., 2001) stably expressing Flag-Merlin were analysed for their ability to co-IP endogenous proteins in either confluent or very sparse cells, where a minimal number of cell-cell contacts are formed (Figure S1E-D). Cell density had no effect on the interactions with VPRBP, CHD1, TAT-SF1 and the SF3A/B complex (Figure 1G and S1E-F). In clear contrast, the interaction with the PAFC and RTF1 was strongly reduced under confluent conditions (Figure 1G and S1E-F). Conversely, Merlin displays a reduced interaction with AMOT and AMOTL1 under sparse conditions (Figure 1G and S1E-F) in agreement with a previous report (Yi et al., 2011). Thus, cell density can differentially modulate Merlin association with different binding partners, with Merlin’s interaction with the PAFC and AMOT family proteins being inversely regulated.

We note that the ability of the PAFC to interact with Merlin in sparse conditions correlates with a slower electrophoretic migration in SDS-PAGE of LEO1 and PAF1 subunits as well as CHD1 (Figure 1F and S1E-F) that is consistent with phosphorylation. This suggests that signalling originating from cell-cell contacts controls post-translational modification of several Merlin interacting proteins in the nucleus, and at least in the case of the PAFC, this correlates with its ability to interact with Merlin.

### The PAFC is required for growth inhibition by Merlin

To map the region of Merlin mediating these novel interactors we expressed GST-fusion proteins of the N- and C-terminal halves of Merlin containing the FERM domain or α-helical and C-terminal domains respectively (Figure 2A). Merlin associates with the PAFC and CHD1 through its FERM domain whereas the interaction with TAT-SF1 could only be detected with the full length protein suggesting several regions of Merlin are involved in this interaction (Figure 2B). As reported, VPRBP also binds to Merlin’s FERM domain (Li et al., 2010) whereas AMOT proteins interact with the C-terminal half (Yi et al., 2011) (Figure 2B).

**Figure 2.**
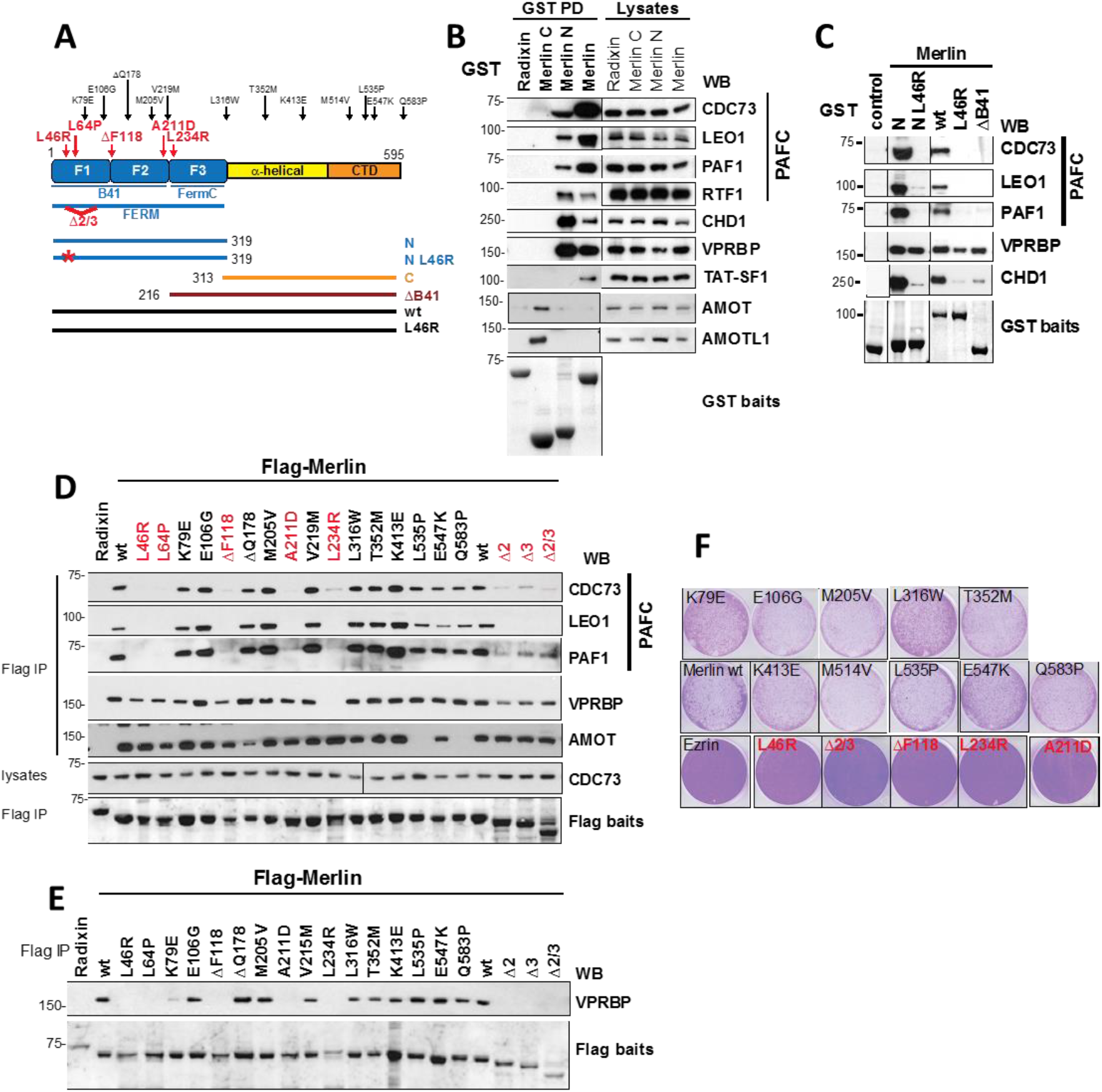
Loss of function mutations in Merlin selectively disrupt the interaction with the PAFC. **(A)** Schematic of Merlin constructs used. **(B)** Merlin’s FERM domain mediates the interaction with the PAFC, CHD1 and VPRBP whereas AMOT and AMOTL1 bind to Merlin’s C-terminus. TAT-SF1 only associates with full length Merlin. GST-Merlin and - Radixin were transfected and purified from HEK293T cells and bound proteins detected by wb. **(C)** The PAFC, CHD1 and VPRBP bind to different regions of Merlin’s FERM domain. Recombinant GST - proteins were used in ‘fishing’ experiments from cell lysates and associated proteins detected by wb. **(D)** Mutations in Merlin selectively disrupt interactions. Flag-Merlin was immunoprecipitated from HEK293T cells and endogenous proteins detected by wb. **(E)** B41-domain Merlin mutants are defective for VPRBP binding under more stringent experimental conditions. As in E but cells were lysed and IPs washed in RIPA buffer containing 0.5% DOC, 0.1% SDS. **(F)** Inhibition of cell proliferation by Merlin correlates with its ability to interact with the PAFC. Clonogenic assays of SF Merlin-deficient SF1335 meningioma cells infected with retroviruses expressing either Ezrin, Merlin wt or Merlin mutants and stained with crystal violet.

The FERM domain is composed of three structurally distinct modules (F1, F2 and F3) that together form a clover-shaped structure. In ‘fishing’ experiments with recombinant proteins, both wild type Merlin and the FERM domain pulled down the PAFC, CHD1 and VPRBP from cell lysates (Figure 2C). The tumour-derived L46R substitution that maps to the F1 lobe disrupted the interaction with the PAFC and CHD1 but only had a partial effect on VPRBP. Similarly, a Merlin deletion mutant lacking the F1 and F2 subdomains of the FERM domain (ΔB41) was unable to interact with the PAFC and CHD1 but could still bind to VPRBP (Figure 2C). To further dissect the interactions, a panel of tumour-derived Merlin mutations distributed throughout the protein were tested for their ability to co-IP endogenous proteins. Mutations within F1 and F2 lobes of the FERM domain (L46R, L64P, ΔF118, A211D) or deletion of exons 2 and 3 within the F1 subdomain disrupt Merlin’s interaction with the PAFC but not VPRBP or AMOT (Figure 2D). The L234R substitution, which lies adjacent to the F2 in the F3 subdomain, does disrupt the interaction with VPRBP (as well as PAFC) (Figure 2D). L535P and Q583P in the C-term domain specifically disrupt interaction with AMOT consistent with its interaction with the C-terminus of Merlin (Yi et al., 2011).

Our findings on the ability of some FERM domain point mutants such as L46R to interact with VPRBP are in contrast to those of Li et al (Li et al., 2010). We reasoned that different experimental conditions could account for these discrepancies since more stringent lysis and washing conditions (RIPA buffer containing 0.1% SDS) were used in their assays. When similar experiments were performed using RIPA buffer, Merlin’s interaction with PAFC, TAT-SF1 and CHD1 can no longer be detected (data not shown). In contrast the Merlin-VPRBP interaction, can still be detected and disruption by FERM domain mutations can now be observed (Fig. 2E) in agreement with Li et al (Li et al., 2010).

Taken together our data suggest that Merlin uses distinct regions of the FERM domain to interact with different proteins, with the F1 and F2 subdomains (B41 domain) mediating the interaction with the PAFC and CHD1 and the F3/FERM-C subdomain primarily mediating the interaction with VPRBP, although substitutions within the other FERM lobes may weaken the affinity for VPRBP and make it more prone to dissociation under certain experimental conditions.

We next assessed the ability of our panel of tumour-derived Merlin mutants to inhibit proliferation when expressed in Merlin-deficient cells SF1335 meningioma cells, where both Merlin isoforms 1 and 2 potently inhibit proliferation in clonogenic assays (Figure 2F). Surprisingly, many of the tumour-identified mutants were still able to inhibit proliferation as efficiently as wild type Merlin (Figure 2F) suggesting these mutations may impair function via decreased protein stability rather than functional activity (Yang et al., 2011) a property that could be overcome by overexpression in our experimental system. Strikingly, only the mutations within the FERM domain (L46R, ΔF118, A211D, L234R) that disrupt the Merlin-PAFC interaction (Figure 2D), were unable to inhibit proliferation of SF1335 cells (Figure 2F). Splice site mutations are frequently found in *NF2* and can lead to deletion of exons 2 and/or 3 (Sainz et al., 1994). A Merlin deletion mutant lacking amino acids coding for exons 2 and 3 was also defective in its ability to inhibit tumour cell proliferation (Figure 2F). Therefore, there is an excellent correlation between the ability of Merlin to inhibit proliferation and its interaction with the PAFC suggesting the Merlin-PAFC interaction is being targeted by inactivating mutations in NF2 tumours.

To study the contribution of the PAFC to growth inhibition by Merlin, Merlin-deficient SF1335 meningioma cells were infected with retroviruses expressing YFP-Ezrin or -Merlin and the cell cycle profile of YFP positive cells was analysed using flow cytometry. Merlin expression led to a pronounced G1 arrest two days after infection whereas expression of Ezrin had no effect (Figure 3A). Knock-down of CDC73 with two different siRNA oligos significantly inhibited Merlin-induced G1 arrest (Figure 3B-D). This strongly suggests that CDC73 is required for Merlin’s tumour suppressor function.

**Figure 3.**
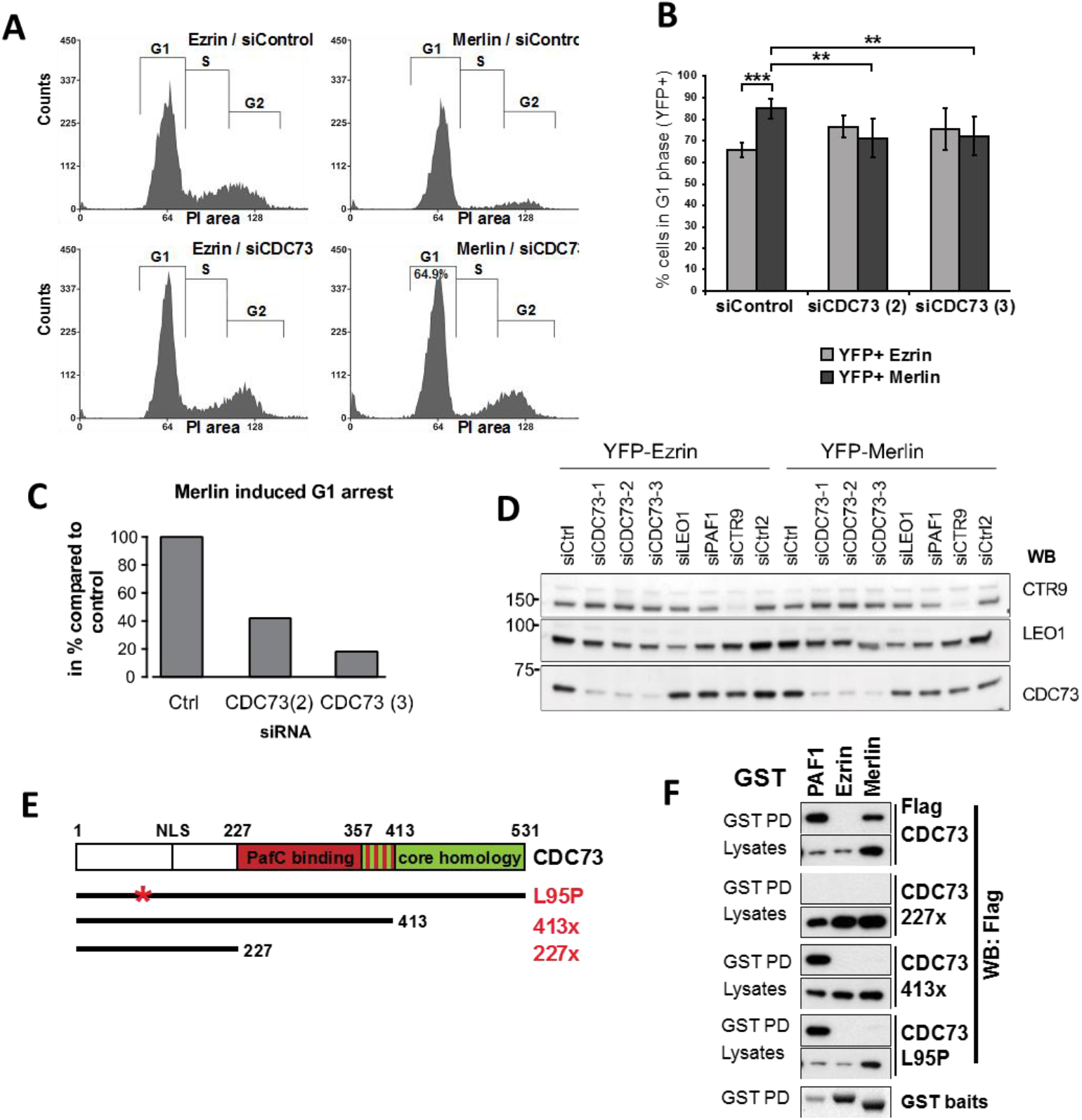
The PAFC is required for growth inhibition by Merlin and the CDC73 and RTF1 subunits are targets of VPRBP-mediated ubiquitination. (A) Knockdown of CDC73 reduces Merlin-induced G1 arrest. SF1335 cells transfected with the indicated siRNAs were infected 24h later with YFP-Ezrin or -Merlin and cell cycle profiles analysed by FACS 48 hours later. Representative experiment of data shown in B. (B) Quantification of experiments as in A. Data represent the mean ±SEM of 5 independent experiments. (*p<0.05, **p<0.01, *** p<0.001) (C) Rescue of Merlin induced G1 arrest by CDC73 knockdown compared to control. One representative experiment shown. (D) Confirmation of CDC73 knockdown by three different siRNA oligos. Western blot analysis of protein lysates from SF1335 cells used in C confirming CDC73 knockdown. (E) Schematic of CDC73 mutants used. (F) Loss-of-function mutations in CDC73 disrupt the interaction with Merlin. GST-PAF1, -Ezrin or -Merlin were co-transfected with Flag-CDC73. Glutathione pull-downs were probed with Flag antibody.

We next considered whether tumour derived inactivating mutations in *CDC73* might exhibit deficiencies in its interaction with Merlin in co-transfection experiments. In contrast with the CDC73 227x truncation mutants that is unable to interact with Merlin or PAFC subunits, CDC73 413x is unable to bind to Merlin (Figure 3F) but still interacts with other PAFC subunits as previously reported (Rozenblatt-Rosen et al., 2005). Similarly, the CDC73 L95P missense substitution (Bradley et al., 2006; Panicker et al., 2010) had no effect on its interaction with the other PAFC subunits but disrupted the interaction with Merlin (Figure 3F). Thus, inactivating mutations in CDC73 disrupt the interaction with Merlin without necessarily influencing the interaction with other PAFC subunits, suggesting that the ability to interact with Merlin is important for CDC73 tumour suppressor function.

### CDC73 is a potential substrate of CRL4^VPRBP^

Merlin interaction with VPRBP E3 ligase complex (CRL4^VPRBP^) in the nucleus is an important mediator of Merlin effects on gene expression (Li et al., 2010). Because VPBRB and PAFC primarily interact with different FERM subdomains, they would be expected to be in close proximity within a complex with Merlin. We thus tested whether subunits of the PAFC could be targets of ubiquitination by CRL4^VPRBP^, as has been described for LATS kinases, themselves Merlin interactors (Li et al., 2014; Yin et al., 2013).

To examine whether CDC73 is ubiquitylated in vivo, HEK293T cells were transfected with HA-ubiquitin and Flag-baits with or without VPRBP. HA blots of Flag IPs showed the laddering pattern consistent with polyubiquitination on Flag-CDC73 but not on other Merlin interactors tested in parallel, such as AMOT and PAF1. Expression of VPRBP leads to a further increase in polyubiquitination associated with CDC73 but not of PAF1 or AMOT (Figure 4A). To test if CDC73 is a direct target of ubiquitylation, similar experiments were performed under denaturing conditions using TAP6-CDC73. Cells were lysed in a guanidinium chloride-containing buffer and subjected to His-tag pulldown with talon beads (Figure 4B). Ub-associated laddering was readily observed on CDC73 purified under denaturing conditions that was further increased by VPRBP, consistent with CDC73 being a direct target of the CRL4^VPRBP^ complex. This effect appears specific to the VPRBP/DCAF1 as expression of other DCAFs had little effect on CDC73 ubiquitination under similar conditions (Figure 4C). To test whether Merlin regulates VPRBP’s ability to ubiquitilate CDC73, similar experiments were performed over-expressing either Merlin or Ezrin as a control. Merlin expression inhibited, albeit modestly, VPRBP stimulated ubiquitination of CDC73 (Figure 4D) consistent with Merlin’s inhibition of CRL4^VPRBP^ activity (Li et al., 2010).

**Figure 4.**
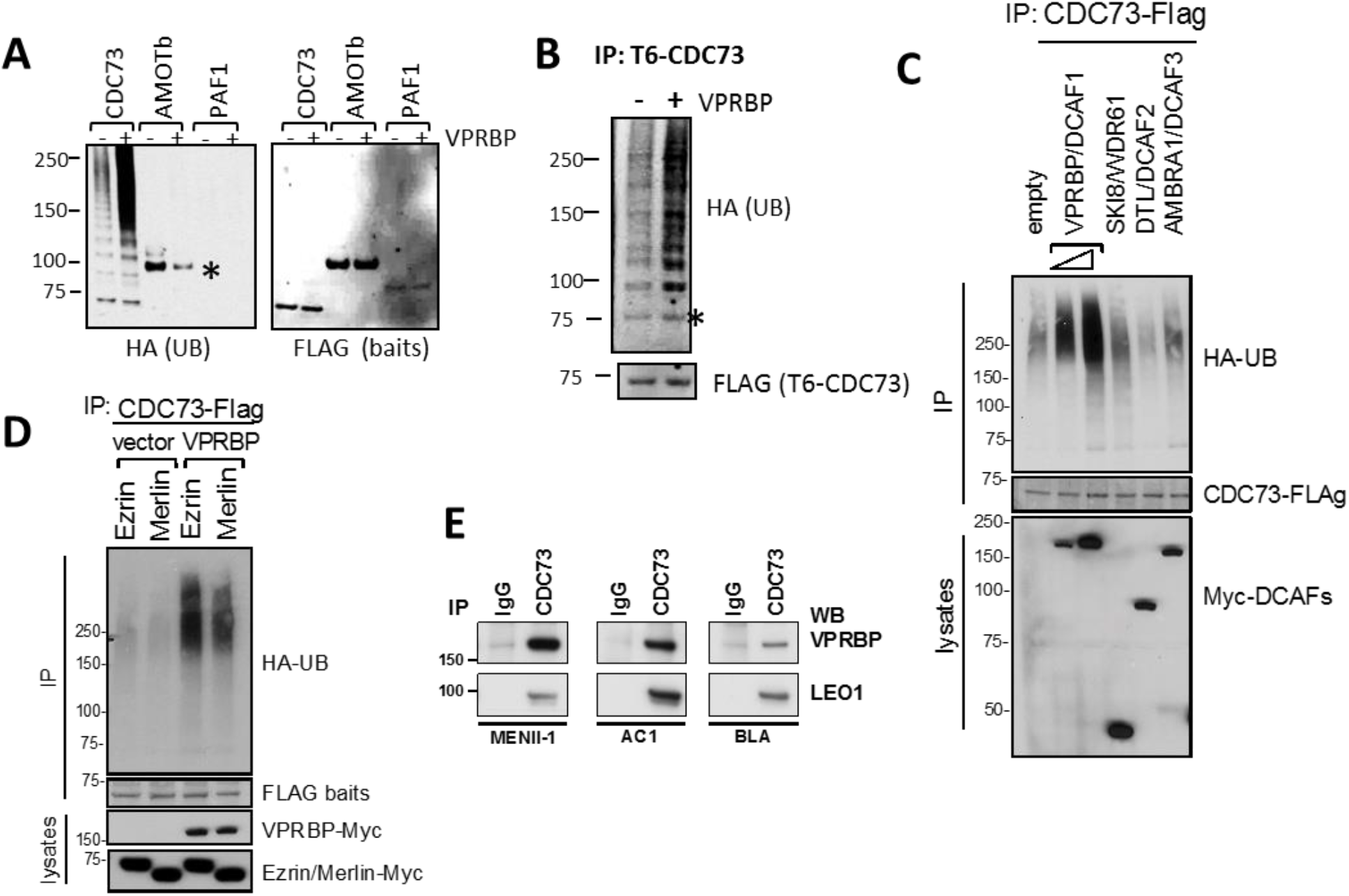
CDC73 interacts with VPRBP and is ubiquitylated in vivo. (A) 293T cells were transfected with HA-ubiquitin, indicated Flag-proteins and either empty vector or VPRBP. Flag IPs were immunoblotted as indicated. The asterisk points to a band corresponding to Flag-AMOTb bait that crossreacts with the HA-antibody. (B) 293T cells were transfected as above with but using TAP6-CDC73 (containing a tandem His tag and Flag tag) and lysed and purified under denaturing condition. Talon beads pull-downs were immunoblotted as indicated. Asterisk points to a band corresponding to the TAP6-CDC73 bait that cross-reacts with the HA antibody on lycor. (C) VPRBP/DCAF1 but not other DCAFs stimulate CDC73 ubiquitination. As in A but comparing Myc-VPRPB to other Myc-DCAFs. (D) Merlin partially inhibits VRPBP stimulated ubiquitylation of CDC73. Flag-CDC73 was transfected with HA-UB and either Merlin or Ezrin as control as in A. Ubiquitin associated with Flag-IPs was detected by wb with an HA antibody (E) Endogenous CDC73 co-IPs with VPRBP in multiple cell lines. WB: Western blot, IP: Immunoprecipitation, PD: Pull down.

VPRBP stimulated ubiquitination of CDC73 correlates with their ability to form a complex *in vivo* as endogenous CDC73 and VPRBP co-IP in multiple cell lines (Figure 4E).Taken together, these observations suggest that CDC73 is a target of CRL4^VPRBP^ mediated ubiquitination and thus Merlin regulation and/or co-recruitment of CRL4^VPRBP^ may provide a molecular mechanism of PAFC regulation by Merlin.

### A gene expression program associated with hypersensitivity to growth inhibition by Merlin

In order to study gene regulation by Merlin, we first characterized expression of Merlin and its interactors in cell lines derived from multiple tumour types. All Merlin interactors tested were found to be ubiquitously expressed (Figure 5A). On the other hand, in addition to mesotheliomas and meningiomas, Merlin expression was absent or aberrant in cell lines derived from diverse tumour types (Figure 5A and S2A-B). Whereas Merlin expression was not detectable in some cell lines, a significant proportion of them (5 out of 11) displayed a faint band of a shorter product in whole cell lysates (Figure 5A) or by IP using two different Merlin antibodies (Figure S2B). This is consistent with splicing mutations as a mechanism of Merlin inactivation (Ahronowitz et al., 2007).

**Figure 5.**
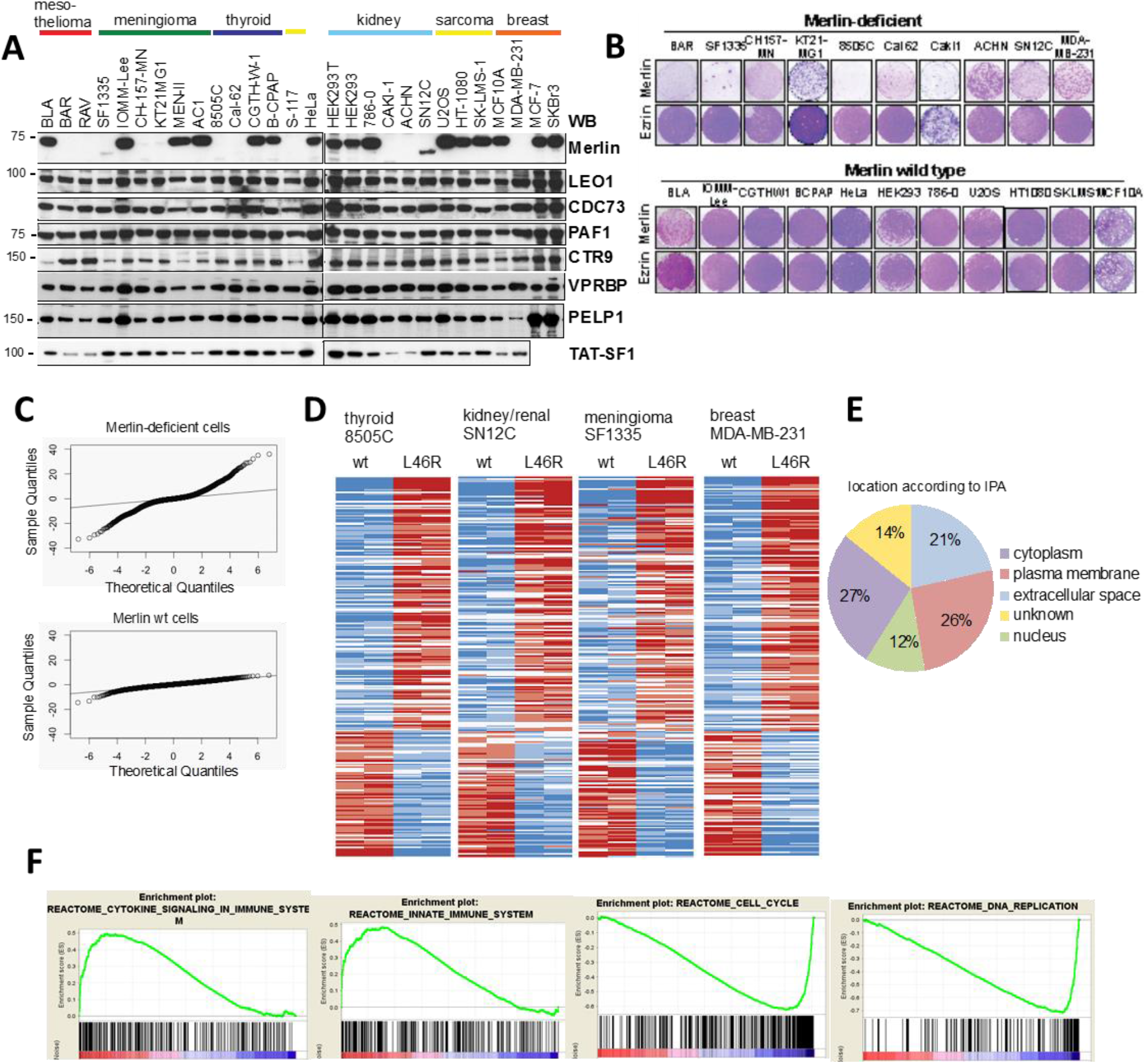
Growth arrest by Merlin is associated with a cross-tissue gene expression program. (A) Merlin protein expression (but not identified interactors) is lost in several tumour cell lines. (B) Merlin-deficient cell lines are hypersensitive to ectopic expression of Merlin. Cells were infected with retroviruses expressing either Ezrin or Merlin and stained with crystal violet 6-12 days later. (C) Merlin expression is associated with far more gene expression changes in Merlin-deficient compared to wild type cells as evidenced by a far greater dispersion of limma t-values compared to the theoretical null distribution (indicated by the black line). (D) Heatmap representing significantly regulated genes in response to expression of wild type Merlin compared to Merlin L46R across four Merlin-deficient cell lines. Blue and red denote low and high expression respectively. (E) Merlin re-expression/hypersensitivity gene signature is enriched in proteins that locate to the plasma membrane and extracellular space. Protein location based on the Ingenuity pathway analysis software IPA (Qiagen). (F) Selected biological pathways from Reactome database regulated by Merlin identified by GSEA. Two representative enrichment plots are shown for up- and down-regulated pathways. See also Figure S4

When Merlin was expressed ectopically, cell lines with wild type Merlin were insensitive to Merlin expression (with only one exception (1 of 11)) and grew at the same rate as Ezrin-expressing or empty vector controls (Figure 5B). In contrast, proliferation of all Merlin-deficient cells (11 of 11 lines) was strongly inhibited by Merlin re-expression (Figure 5B). This is consistent with the concept of ‘oncogene addiction/tumour suppressor gene hypersensitivity’ (Weinstein, 2002) and suggests that in tumours with *NF2* mutations, absence of Merlin expression is required for proliferation, regardless of tumour type and/or of additional somatic mutations (Figure S2A).

To study how Merlin selectively induced growth arrest in Merlin-deficient cells and shed light on Merlin-regulated genes, we analysed by microarray global gene expression after expressing either Merlin wt or an inactive L46R mutant that is unable to interact with the PAFC (Figure 2D) in a set of four cell line pairs derived from different tumour types (Figure S2C). Whereas in Merlin-deficient cells, re-expression of wt Merlin causes large changes in gene expression, the response in Merlin wt cells is marginal (Figure 5C), with only 5 significant changes (fold change >1.5, q<0.05) (Figure S2D and Table S1). In contrast, there was a consistent cross tissue signature of 247 gene changes (85 down, 162 up, fold change >1.5,q<0.05) shared across the 4 different Merlin-deficient cell lines (Figure 5D and Table S2) hereby referred to as the Merlin reexpression/hypersensitivity signature. Therefore, the ability of Merlin to suppress proliferation correlates with its ability to regulate a characteristic gene expression program. Validation of selected Merlin-regulated genes by quantitative RT-PCR is shown in Figure S3.

Ingenuity pathway analysis revealed the Merlin gene expression signature is highly enriched for genes encoding plasma membrane and extracellular proteins (Figure 5E) suggesting a role for Merlin in cellular communication with the microenvironment. Knowledge based pathway analysis by gene set enrichment analysis (GSEA) (Subramanian et al., 2005) revealed Merlin re-expression caused downregulation of genes involved in cell cycle progression and DNA replication (Figure 5E and S4) consistent with the growth inhibition observed in Merlin-deficient cells. On the other hand, upregulated pathways suggest a transcriptional role for Merlin in inflammation and innate immunity (Figure 5E and S4). Consistent with a functional link between Merlin and Hippo signaling a conserved YAP transcriptional signature is strongly downregulated by Merlin expression (Figure S4C). Coordinated up/down regulation of several transcriptional signatures also suggests possible functional relationships between Merlin and other transcriptional pathways (Figure S4C).

### Merlin is involved in the transcriptional response to high cell density

To further define genes regulated by Merlin and their relationship to contact inhibition, microarray experiments were performed after Merlin depletion in HMLE cells at both high and low cell densities. HMLE cells where Merlin expression was inhibited show far fewer significant changes than control cells in response to high cell density (Figure S5A): In control cells 801 (449 up, 352 down) genes were significantly regulated (fold change >1.5, p<0.05) compared to only 60 (48 up, 12 down) in Merlin knock-down cells (Figure S5B and Tables S3 and S4). Validation by qPCR of selected genes regulated by high cell density is shown in Figure S6C.

Comparison of a high cell density gene signature with the Merlin reexpression/hypersensitivity signature shows a significant positive overlap (chi-square, p < 0.001, Figure SED). Out of the 251 genes regulated by Merlin re-expression in tumour derived cell lines over 20% (52 genes) are regulated by high cell density in the same direction in HMLE cells (Table S5). This is consistent with growth suppression by Merlin being mediated at least partly by some of the same transcriptional changes that mediate contact dependent inhibition of proliferation. Merlin knockdown caused substantially more transcriptional changes in dense cells than in sparse cells consistent with its role in contact inhibition. However, Merlin inhibition in sparse cells still induced substantial changes suggesting Merlin has a wider role in transcription beyond regulation by high cell density (Figure S5E, Tables S6 and S7).

To verify the relevance of these gene signatures, we used the Oncomine concepts platform (https://www.oncomine.org). Genes that are negatively regulated by Merlin in our signatures overlap significantly with overexpressed genes in NF2-deficient cells from a multi cancer cell line study (Garnett et al., 2012) (Figure S5G) validating the integrity of our signatures. We reasoned that regulated genes moving in the opposite direction in the re-expression/hypersensitivity and knockdown signatures would meet the most stringent criteria for Merlin target genes and are hereby defined as the Merlin core signature (Figure S5H and Table S12). Strikingly, a majority (~ 63%) of the genes within this signature code for cell surface-bound and secreted proteins (Figure S5H) highlighting again a key role for Merlin’s transcriptional response in cell communication with the microenvironment.

### Merlin regulates PAFC association with the chromatin of a subset of genes

Given Merlin’s physical interaction with the PAFC, we next set out to study whether Merlin can regulate the association of PAFC with the chromatin of selected genes upon re-expression in Merlin-deficient cells. In an initial study using EGF-stimulated association with FOS gene by chromatin immunoprecipitation followed by PCR (ChIP-PCR) (Chen et al., 2009) we found that an antibody against the LEO1 subunit of the PAFC gave considerably higher signal-to-noise ratios than the antibodies against CDC73, PAF1 or CTR9 subunits tested (data not shown). Similarly, among the Merlin-deficient cells tested, MDA-MB-231 gave the highest signal to noise ratios. Therefore a LEO1 antibody was used, in parallel with a RNA pol II antibody, for ChIP-followed by massive parallel DNA sequencing (Chip-seq) in MDA-MB-231 cells infected with lentiviruses expressing either Merlin wt or the PAFC interaction-defective L46R Merlin mutant.

The vast majority of RNA Pol II bound genes are also bound by LEO1 (Figure 6A), in line with a general role of the PAFC in gene expression (Jaehning, 2010). When the binding profiles for representative genes are compared, RNA Pol II is found predominantly at the transcription start site (TSS), whereas LEO1 peaks downstream of the TSS and follows RNA Pol II throughout the coding region (Figure 6B), consistent with a role of the PAFC in elongation (Jaehning, 2010).

**Figure 6.**
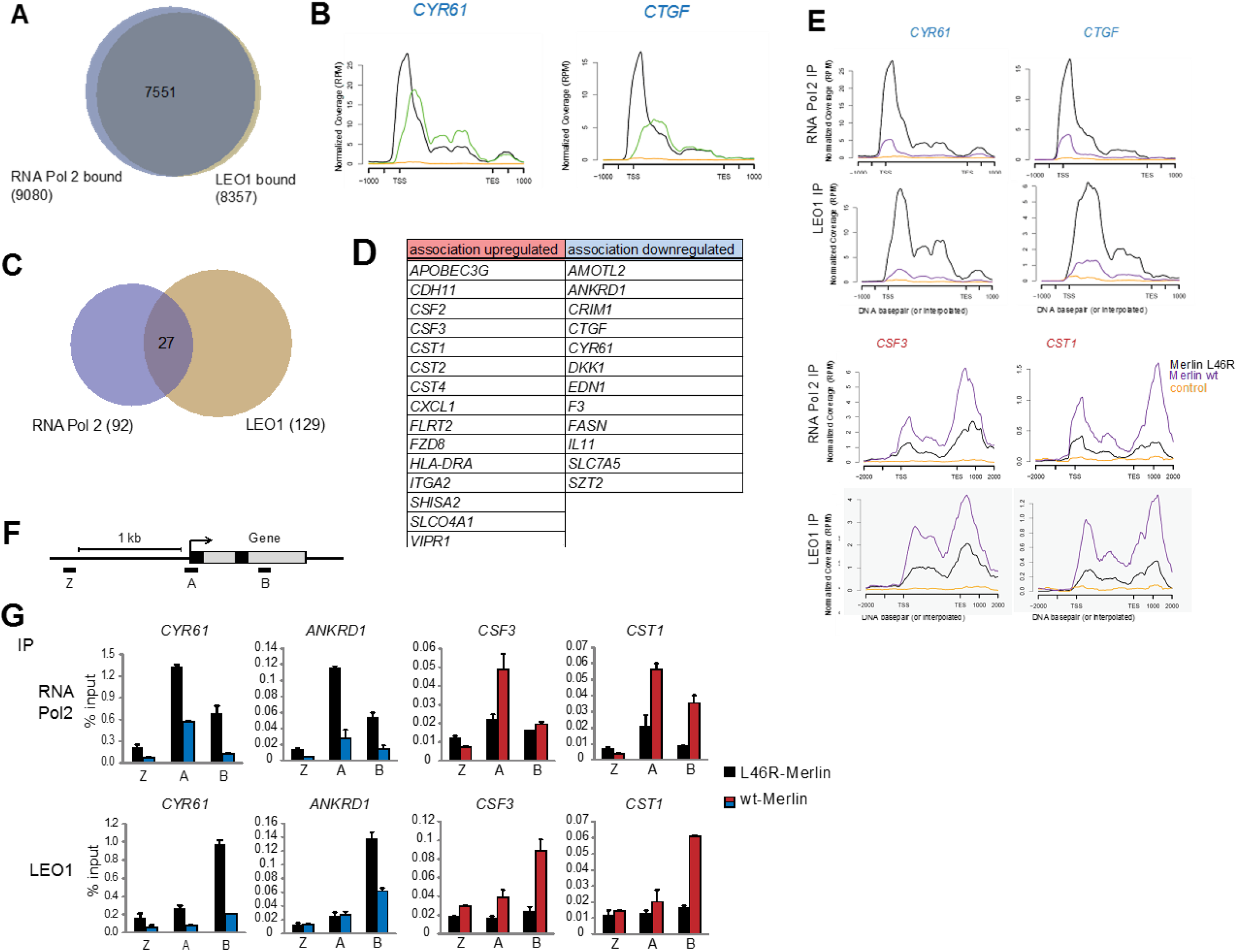
Merlin regulates PAFC association with chromatin of a subset of genes. **(A)** Most RNA Pol II bound genes are also bound by LEO1. Overlap of RNA Pol II- and LEO1-bound genes in Merlin L46R cells as determined by ChIP-seq in MDA-MB-231 cells. **(B)** Plots of the short read coverage across the genomic regions of CYR61 and CTGF example genes showing LEO1 (green) association peaks downstream of RNA Pol II (black). Antibody control is in orange. **(C)** Overlap of RNA Pol II and LEO1 bound genes significantly regulated by wild type Merlin expression (>1.5-fold change, >2-fold over ctrl antibody IP). **(D)** Genes from C where both RNA Pol II and LEO1 association is regulated by Merlin expression. **(E)** Plots of the short read coverage for selected genes showing regulation of RNA Pol II and LEO1 association by wt (purple) compared to mutant (black) Merlin expression. Blue and red denote down- and up-regulation by Merlin expression. **(F)** Schematic showing location of primers used in G. **(G)** Validation of ChIP-seq results. Chromatin from MDA-MB-231 cells expressing wild type or L46R Merlin was IPed using a RNA Pol II, LEO1 and control antibody and qPCRs performed with the indicated primers. Error bars represent SD from two independent PCRs. Representative of three independent experiments. TSS: transcriptional start site, TES: transcriptional end site.

Expression of wt Merlin regulates the association of LEO1 and RNA Pol II with only a relatively small subset of genes (129 on LEO ChiP, 92 on Pol II ChIP, fold change >+/− 1.5) (Figure 6C, Tables S9 and S10). With the cut offs used, 27 genes were identified where Merlin regulates both RNA Pol II and LEO1 binding (Figure 6D). The majority of these (20 of 27) were also found regulated by Merlin in the microarray experiment in MDA-MB-231 cells. Plots of the short read coverage for selected Merlin-regulated genes are shown in Figure 6E and validation by ChIP-qPCR in Figure 6G. Taken together these results suggest that Merlin can regulate the ability of the PAFC to associate with selected genes and this correlates with the concomitant regulation of their transcript levels.

### Proteomic analysis of the CDC73 interactome highlights a key role for FAT cadherins in Merlin’s interaction with the PAFC

In an effort to better understand at the molecular level how Merlin regulates PAFC function we used affinity proteomics to investigate the PAFC interactome and its regulation by Merlin. TAP6-tagged PAF1, LEO1 and CDC73 subunits of the PAFC were stably expressed in Merlin wild-type HT1080 and Merlin-deficient MDA-MB-231 cells. Affinity purification followed by mass spectrometry (AP-MS) identified in all cases the other subunits of the PAFC as the main co-purifying proteins. Other proteins were also co-purified, mostly with roles in RNA processing and transcription although surprisingly there also was also a cluster of proteins associated with cell-cell contacts (Figure S6A and Table S11). Many co-purified with all 3 subunits consistent with copurification of the whole PAFC complex, but subunit-specific as well as cell line-specific interactors were also found (e.g. FAT1 and TP53 in Merlin-deficient MDA-MB-231 cells and FTSJ3 and ZZEF1 in Merlin wt HT1080 cells (Figure S6B-C, Table S11).

To further investigate Merlin regulation of the PAFC interactome while avoiding variability associated to inter-cell line comparisons, we performed AP-MS of the TAP6-CDC73 subunit in HEK293T cells co-expressing either Merlin or Ezrin as a control. Most purifying proteins identified were nuclear proteins involved in transcription. However, there also was a significant fraction of proteins associated with cell junctions and cell adhesion (Figure 7A and Table S12) suggesting a direct link between cell-cell contacts and CDC73 function. Using peptide counts as a semi-quantitative measurement, Merlin expression decreased association with several proteins, mainly involved in cell adhesion, whereas it increased association of other proteins mainly involved in transcription and chromatin remodeling, including subunits of the Nucleosome Remodeling Deacetylase (NuRD) complex (Figure 7B and Table S12). Therefore, Merlin can both positively and negatively regulate CDC73 association with different sets of proteins.

**Figure 7.**
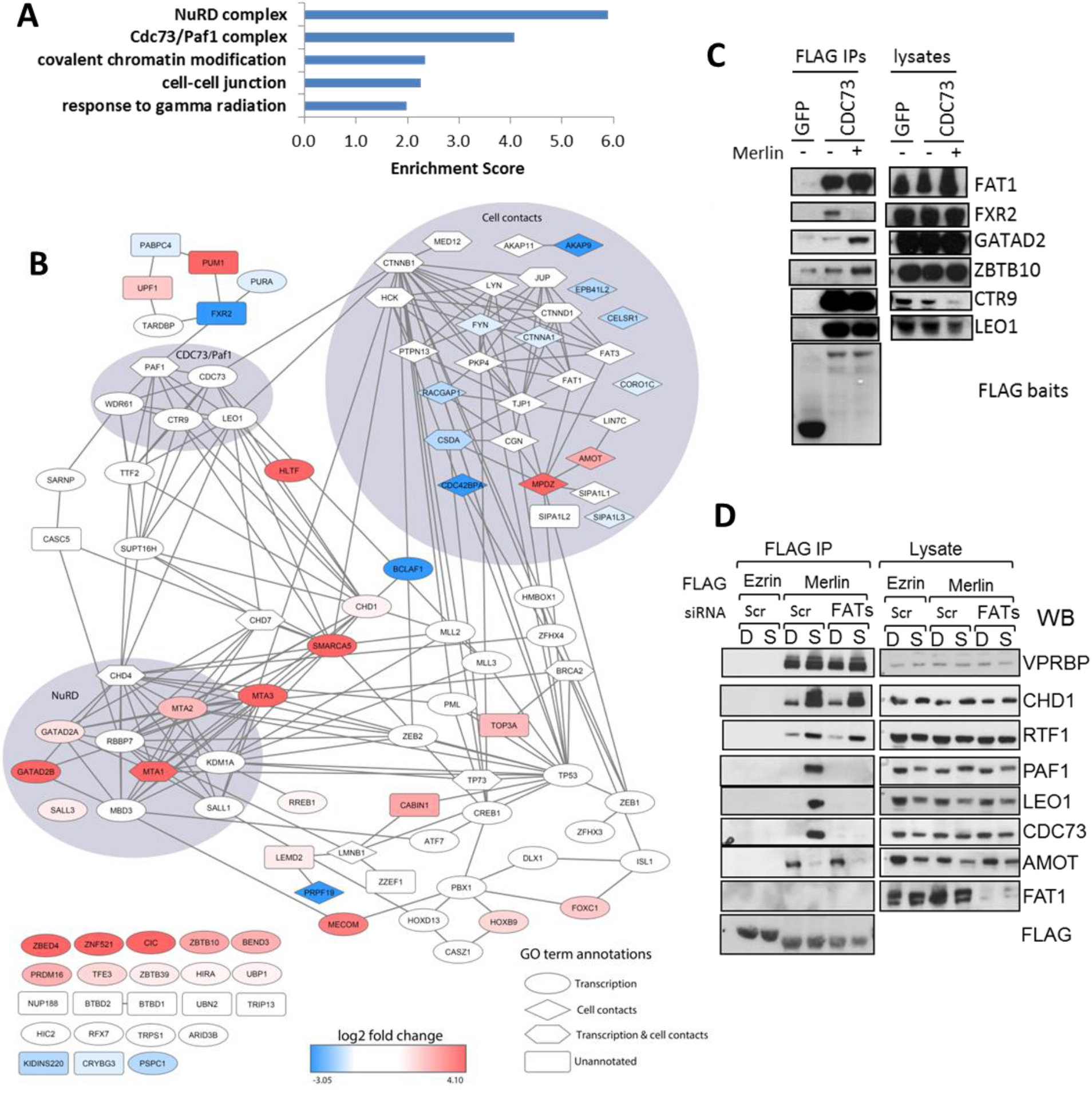
FAT cadherins interact with CDC73 and are necessary for Merlin interaction with PAFC. (A) Gene Ontology term enrichment of the CDC73 interactome in 293T cells. TAP6-CDC73 was affinity purified from 293T cells co-expressing either Merlin or Ezrin. Co-purifying proteins were subjected to functional annotation analysis with GO biological process, cellular component and molecular function terms using DAVID (Huang da et al., 2009). Top 5 results shown. Terms correspond to GO:0010332, GO:0005911, GO:0016569, GO:0016593 and GO:0016581. (B) Bioinformatics analysis of the CDC73 interactome and its regulation by Merlin. Proteins from the STRING functional network were annotated with log2 fold change regulation in Merlin-vs Ezrin-expressing cells (red, upregulated; blue, downregulated in cells over-expressing Merlin; white, unchanged). The shape of protein nodes represents the functional annotations (ellipse: transcription; diamond: cell contacts; hexagon: annotated with both transcription and cell contacts; rectangle: not annotated with these terms). Proteins highlighted with a background surface were annotated with functional terms and cluster together in the network. See also Table S12 and supplementary methods. (C) Validation of some interactions identified by CDC73 AP-MS. Flag-CDC73 or Flag-GFP were expressed in 293T cells with either empty vector or Merlin and Flag-IPs probed with the indicated antibodies. (D) FAT cadherins are required for Merlin interaction with PAFC. 293T cells were transfected with scramble or siRNAs against FAT1, FAT3 and FAT4 and 24 hours later transfected with either Flag-Ezrin or Merlin. Flag immunoprecipitates from dense or sparse cells were probed as indicated.

A comprehensive validation of all these candidate direct or indirect interactions is beyond the scope of this study. However, we were particularly intrigued by the link between CDC73 and cellcell adhesion and in particular with its association with FAT cadherins. FAT1, which like CDC73 functions as a tumour suppressor (Morris et al., 2013), was found associated with CDC73 by AP-MS in both 293T and MDA-MB-231 cells (Table S11 and S12) and we thus focused on this interaction for this study. Endogenous FAT1 can be readily detected associating with Flag-CDC73 in co-IP experiments and this interaction is not affected by Merlin expression (Figure 7C), consistent with the proteomics data in 293T cells. This, together with the observation that FAT1 also copurified with CDC73 in Merlin-deficient MDA-MB-231 cells (Table S11), suggests CDC73 and FAT 1 interact independently of Merlin.

We next investigated whether FAT proteins could regulate the Merlin-PAFC interaction. We used siRNAs to knock-down all FAT isoforms expressed in 293T cells (FAT1, FAT3 and FAT4 as determined by RT-PCR, data not shown) and assessed Flag-Merlin interactions with endogenous proteins in both dense and sparse cells as before (see Figure 1 and S1). Strikingly, knock-down of FAT cadherins strongly inhibits Merlin interaction with PAF1, LEO1 and CDC73 PAFC subunits under sparse conditions whereas there was no effect on other Merlin interactors such as VPRBP, AMOT, CHD1 or RTF1. Thus, FAT cadherins are required for Merlin’s association with the PAFC.

## DISCUSSION

We show that Merlin interacts with a set of nuclear proteins involved in gene expression at post-initiation steps including the PAFC, RTF1, the CHD1 chromatin remodeller and the TAT-SF1 elongation/splicing factor. Interactions between the PAFC, RTF1, CHD1 and TAT-SF1 have been previously reported and suggest a shared role in elongation and splicing (Chen et al., 2009; Sims et al., 2007).

Several of these Merlin interactors are themselves deregulated in cancer. The CDC73 subunit of the PAFC functions as a tumour suppressor (Newey et al., 2010) and we show CDC73 is required for growth inhibition by Merlin. Furthermore, tumour derived inactivating CDC73 mutations disrupt its interaction with Merlin whereas conversely, inactivating Merlin mutations disrupt the interactions with the PAFC. Furthermore, we show that the interaction between Merlin and the PAFC is regulated by FAT cadherins, which are themselves mutated across various cancer types (Morris et al., 2013; Sadeqzadeh et al., 2014). Additionally, CHD1 is mutated in prostate cancer (Grasso et al., 2012; Huang et al., 2012; Liu et al., 2012) and through TAT-SF1, Merlin interacts with the U2 snRNP complex of the spliceosome, whose components are also frequently mutated in several cancer types (Escobar-Hoyos et al., 2019).Taken together these findings suggest a function for nuclear Merlin as part of a tumour suppressor network that helps regulate expression of target genes at the level of elongation, chromatin remodelling and RNA processing.

Growth inhibition by Merlin is associated with its ability to regulate a cross-tissue gene expression program that is also regulated by cell density. Merlin interacts with α-catenin and Angiomotin junctional proteins and thus is well poised to respond to signals originating from cellcell contacts. Cell density is known to regulate Merlin phosphorylation and activity (Li et al., 2012) and we show that subunits of the PAFC as well as CHD1 are also likely phosphorylated in a cell density dependent manner. Moreover, Merlin’s ability to interact with the PAFC is regulated by FAT cadherins and correlates with the ability of FATs to associate with the CDC73 subunit of the PAFC. Taken together, these results are consistent with a model whereby multiple signals originating from cell-cell contacts converge at the level of Merlin-PAFC interaction to mediate a growth inhibitory gene expression program associated with contact inhibition of proliferation (Figures S7).

Although transmembrane proteins with a large extracellular domain, FATs are processed proteolytically by mechanisms similar to Notch and nuclear localization of FAT1 cytoplasmic tail and association with transcription regulators has been reported (Fanto et al., 2003; Feng and Irvine, 2009; Hou and Sibinga, 2009; Magg et al., 2005; Sadeqzadeh et al., 2011). Interestingly, CDC73/Parafibromin has been reported to interact with the Notch intracellular domain (as well as B-catenin and Gli1) (Kikuchi et al., 2016). Conversely although CDC73 is predominantly nuclear, it has also been detected in the cytoplasm in some contexts (Agarwal et al., 2008; Jo et al., 2014). Future studies should shed further light on the nature of the interaction between the CDC73 and FAT tumour suppressor proteins but their association with renal defects may provide an independent genetic link: in addition to cancer, FAT1 mutations are linked to glomerulotubular nephropathy and fat1 knockdown in zebrafish causes pronephric cysts (Ciani et al., 2003; Gee et al., 2016) whereas CDC73 mutations in HPT-JT syndrome are associated with renal lesions in 20% of cases, most commonly cysts (Jackson et al., 1993).

Elongation control plays a crucial role in regulated gene expression and provides an additional regulatory node in the transcription cycle to exert combinatorial control of transcription levels (Jonkers and Lis, 2015; Nechaev and Adelman, 2011). In contrast to initiation, where target gene specificity can be provided by sequence specific transcription factors, far less is known about how elongation and RNA processing can be differentially regulated, with likely multiple mechanisms involved for individual genes or network of genes (Jonkers and Lis, 2015; Nechaev and Adelman, 2011). The PAFC travels with RNA Pol II on most actively transcribed genes but Merlin can target the PAFC to associate with only a subset of genes. We propose that Merlin may function as a co-factor to target a protein complex involved in elongation and RNA processing to a particular subset of genes regulated by cell adhesion.

The molecular mechanisms involved remain to be determined but CRL4^VPRBP^-mediated ubiquitination is likely to play a role. Merlin independently interacts with both CRL4^VPRBP^ and PAFC and is thus expected to bring them into the same macromolecular complex and we show that the CDC73 subunit of the PAFC is a likely target of CRL4^VPRBP^-mediated ubiquitination. How this modification is regulated *in vivo* and how it affects PAFC function remains to be determined. It is worth noting that VPRBP was originally identified as the HIV VPR interacting protein and that many of the Merlin interactors identified in our study, including TAT-SF1, PAFC and CHD1, are used by HIV during its transcription cycle (Gallastegui et al., 2011; Sobhian et al., 2010; Vanti et al., 2009). Although the role of VPR in the HIV cycle is still unclear, it is tempting to speculate that it may function in a manner equivalent to Merlin to target VPRBP to regulate PAFC-mediated transcription of viral target genes.

The genomics data of this study is consistent with the known role of Merlin as an upstream regulator the Hippo pathway pathway (Cooper and Giancotti, 2014; Ma et al., 2019) as many of the genes in our Merlin signatures are known YAP target genes. Our study suggest that Merlin may provide independent layers of regulatory control in the transcription cycle of some genes (Figure S7): According to our proposed model, Merlin at the membrane primarily regulates LATS phosphorylation of YAP/TAZ to promote TEAD-dependent Pol II recruitment and transcription initiation, whereas nuclear Merlin, through its interaction with the PAFC (as well as CHD1, RTF1, TAT-SF1 and others) may also regulate post-initiation steps such as pause release, elongation rate and/or RNA processing in a subset of target genes (Figure S7).

Understanding these multiple regulatory nodes within the Hippo pathway, how the function of nuclear Merlin is intertwined with its membrane function and how it helps coordinate the different steps of the transcription cycle of selected genes through its interaction with a network of nuclear tumour suppressor proteins will be important to be able to provide new therapeutic opportunities for NF2 tumours as well as the many human cancers where this pathway is deregulated

## EXPERIMENTAL PROCEDURES

### Interaction assays

Cells were lysed in PBS-E LB (1x PBS, 1% TX100, 1 mM EDTA, 1 mM DTT, Protease Inhibitor cocktail (Roche) and Phosphatase Inhibitor solution (Sigma)). When indicated RIPA buffer (PBS-E LB+ 0.1% SDS, 0.5% DOC,) was used. Flag (M2) agarose beads (Sigma-Aldrich), glutathione sepharose beads (GE Healthcare), or a combination of the appropriate antibody and Protein A or G sepharose beads (GE Healthcare) were used to immunoprecipitate tagged or endogenous proteins from cleared lysates. For cell density experiments, when 10cm dishes reached ~90% confluency, either medium was changed (Dense) or split to 4-5×15cm dishes (Sparse) and cells harvested 12-15 hrs later. Immunoprecipitates were extensively washed with lysis buffer minus inhibitors, drained and resuspended in NuPAGE LDS sample buffer (Life Technologies).

### Clonogenic assays

Cells were infected with retroviruses or lentiviruses expressing either Merlin or Ezrin, seeded at different densities in 6-well plates and stained with crystal violet (0.5% in 10% Methanol) 7-14 days later when the Ezrin or empty vector control wells reached confluency.

### Flow-cytometry

Cells were fixed in 70% Ethanol two days after retroviral infection with YFP-Ezrin/Merlin and cell cycle profiles assessed by propidium iodide (PI) staining. The cell cycle of YPF-positive cells is shown, no changes were seen in the YFP-negative population of cells. Where indicated cells were transfected with siRNA, infected 24 hrs later with retroviruses expressing YFP constructs and harvested 48 hrs later for FACS analysis.

### Ubiquitination assays

Cells were co-transfected with pMT123-HA-ubiquitin (Treier et al., 1994) and lysed 2 days later. For denaturing conditions, cells were lysed with Guanidine lysis buffer (6M Guanidine-HCl, 50mM NaHPO4 buffer pH:8, 0.3M NaCl, 0.1% TX-100, 10mM imidazole, 5mM b-Mercaptoethanol), briefly sonicated and incubated after centrifugation with TALON beads. Beads were washed once with Guanidine LB and 3 times with Urea wash buffer (8M urea, 50mM NaHPO4 buffer pH:8, 0.3M NaCl, 0.1% TX-100, 10mM imidazole, 5mM b-Mercaptoethanol). For native conditions, cells were lysed in RIPA buffer (1x PBS, 1% TX100, 0.1% SDS, 0.5% Sodium deoxicholate,1 mM EDTA, 1 mM DTT, with Protease and Phosphatase Inhibitors). Cleared lysates were incubated with Flag beads for 1-2 hours and beads washed 4 times with RIPA LB. Drained beads were resuspended in LDS sample buffer and run in Nupage 4-12% gels. After gel transfer, membranes were probed with anti-HA-peroxidase (Roche-12013819001) and developed by chemiluminescence or with FLAG-Dylight680 and HA-IRDye800 (Rockland) and developed with the Odissey imaging system (Lycor).

## ACKNOLEDGEMENTS

We thank Arnold Pizzey for help and advice with the FACS analysis; Anita Lal, David Gillespie, Julian Downward and Marco Giovannini for cell lines; Richard Jenner for help with ChIP assays, Nic Tapon, Richard Jenner, Paolo Salomoni, Benoit Bilanges and Daniel Hochhauser for critically reading the manuscript. K. Klupsch was funded by a research fellowship from the Deutsche Forschungsgemeinschaft (DFG). The UCSF Mass Spectrometry Facility (A.L. Burlingame, Director) was supported by the Biomedical Research Technology Program of the National Center for Research Resources, NIH NCRR RR001614, and NIH NCRR RR012961.

## SUPPLEMENTARY FIGURES

**Figure S1.**
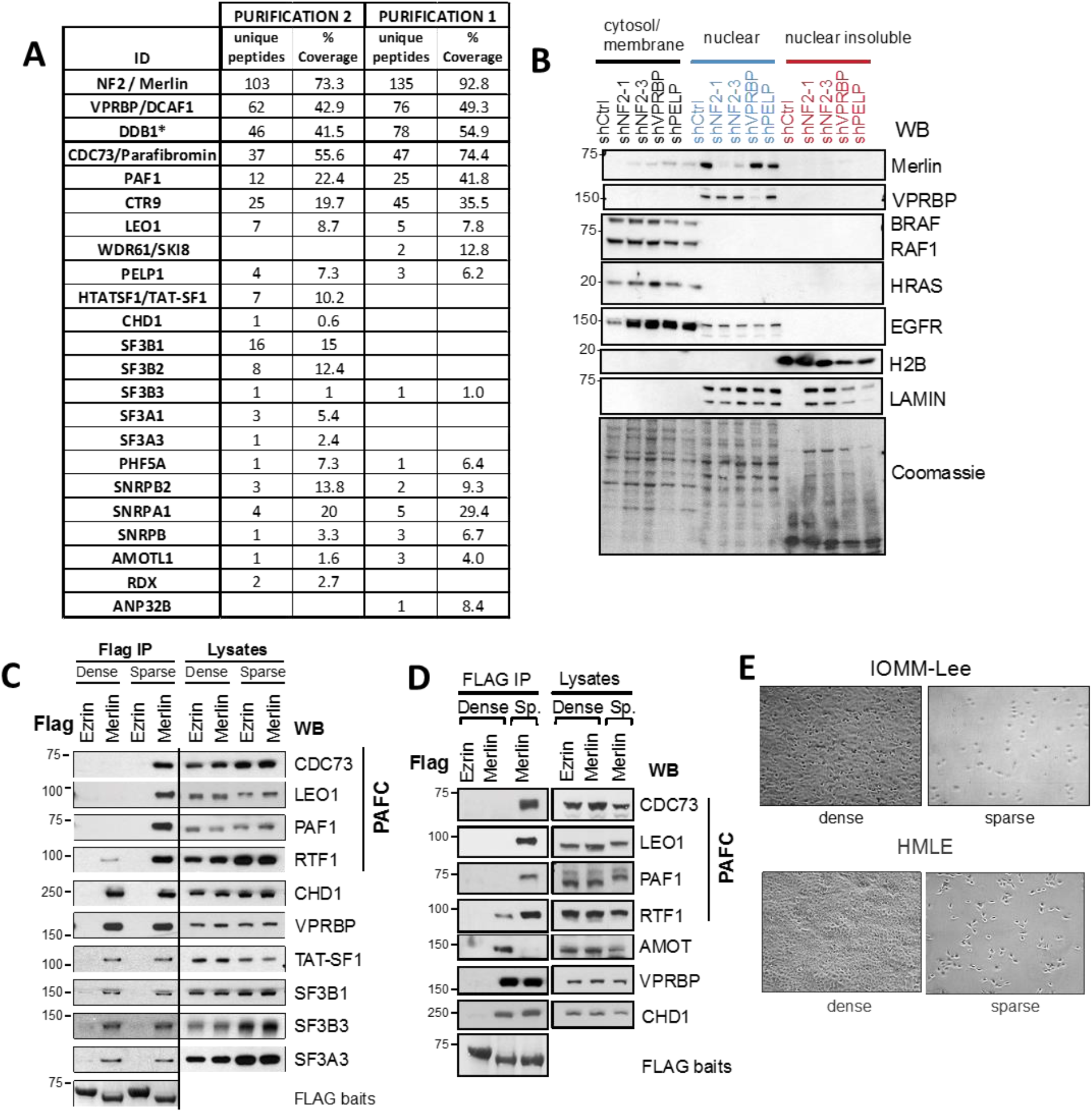
Merlin interacts with the PAFC in a cell density dependent manner. (A) Proteins identified by mass spectrometry after 2 independent Merlin affinity purifications from HEK293T. * Some peptides for no DDB1 were also found in the SHOC2 control bait used. (B) Merlin resides primarily in the nuclear fraction. IOMM-Lee cells expressing various shRNAs were fractionated into a cytoplasmic/membrane, nuclear and nuclear insoluble fractions and proteins detected by Western blot. The shRNA knockdown of Merlin and Vprbp serve as specificity control for the immunoblot. BRAF/CRAF are used as examples of cytosolic proteins whereas HRAS and EGFR are examples of membrane proteins. Some EGFR can also be detected in the nucleus consistent with previous observations (Wang et al., 2010). Histone H2B is a marker for nuclear insoluble fraction. (C) Merlin’s interaction with the PAFC is inhibited in confluent cells. Flag IPs from dense or sparse IOMM-Lee cells stably expressing Flag-Ezrin or Merlin were probed with the indicated antibodies. (D) As in D, but using HMLE cells. (E) Representative images of cell densities at time of the experiments in C and D.

**Figure S2.**
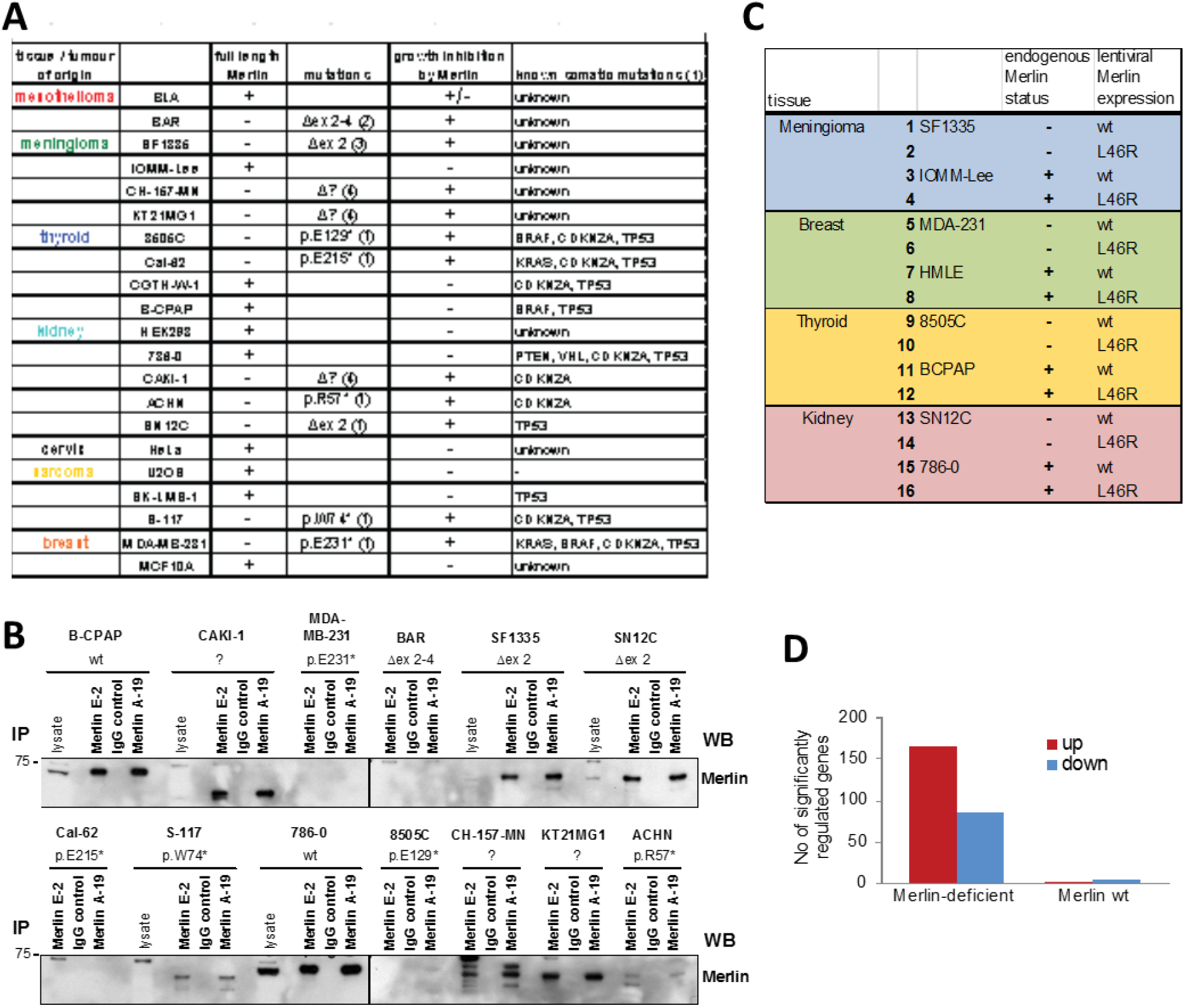
Merlin regulates gene expression in Merlin deficient cells. (A) Summary of proliferation studies for all tested cell lines in Figure 5. Known NF2 mutations as well as other somatic mutations are listed. (1) COSMIC database, (2) (Deguen et al., 1998), (3) M. Giovannini personal communication, (4) see S2B. (B) Characterization of Merlin expression by IP-wb. Merlin was IPed using antibodies directed against the N-terminus (A-19) and the C-terminus (E-2) and detected by wb using an anti-C-terminus antibody (Bethyl). Cal62, 8505C and ACHN show no Merlin protein expression, whereas CAKI-1, SF1335, SN12C, CH-157-MN, KT21MG1 express shorter protein products consistent with splice site mutations. (C) Experimental design of gene expression microarray experiment in tumour cell lines. Merlin deficient and Merlin wild type cell lines from 4 different tissue types were infected with lentiviruses expressing either wild type or L46R Merlin, RNA was isolated 48 hrs later and analysed using GeneChip^®^ Human Gene 2.0 ST Arrays (Affymetrix). (D) Merlin expression regulates gene expression in Merlin deficient but not Merlin wild type cells. Number of significantly regulated genes (q ≤ 0.05, fold change >1.5) identified by microarray.

**Figure S3.**
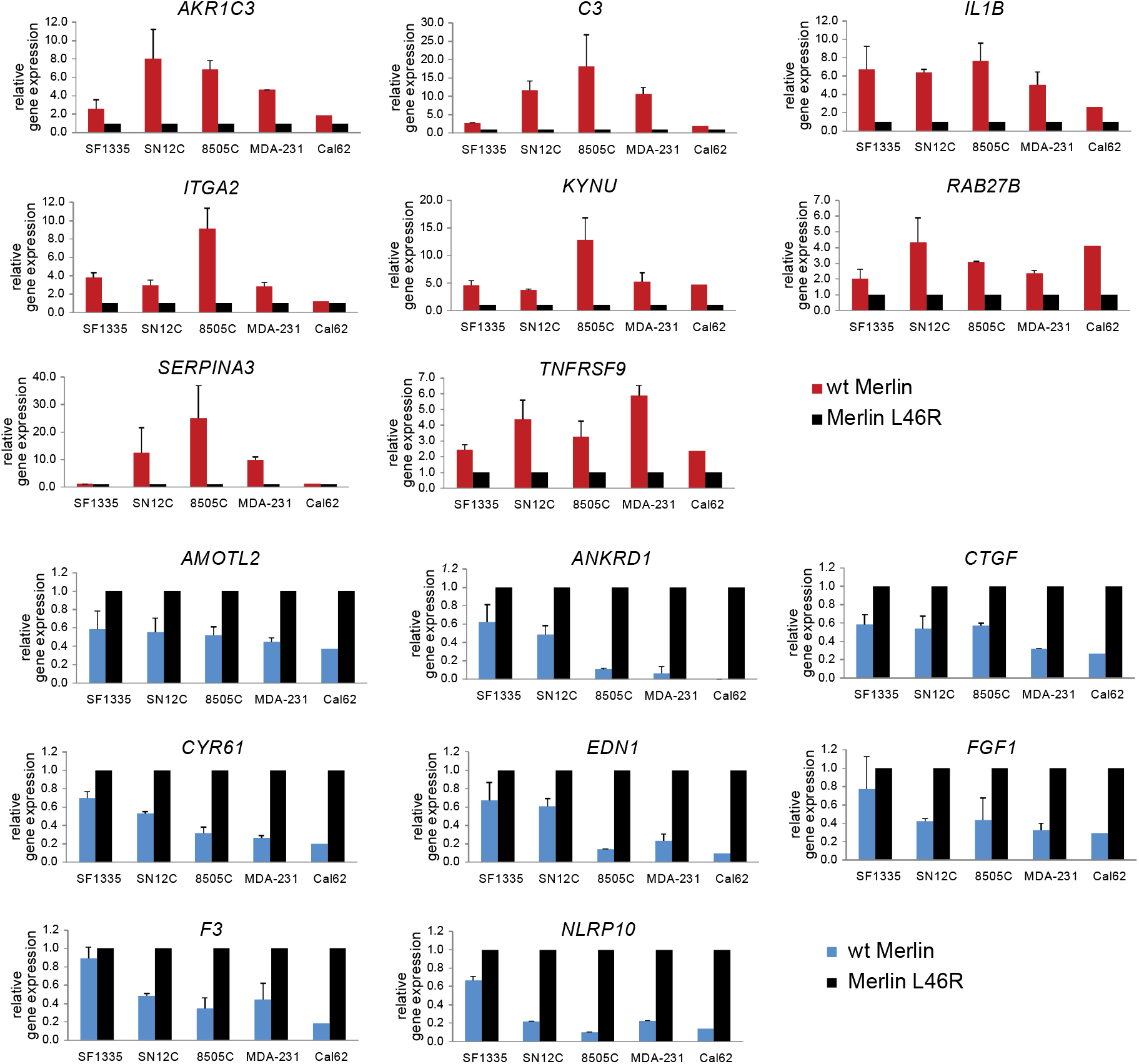
Validation of microarray results shown in Figure 5 by RT-qPCR. cDNA from cells as in S2C was analysed by qPCR. Data is represented as gene expression relative to actin (ACTB) and normalized relative to Merlin L46R mutant. Error bars indicate SD from three independent experiments. Cal62 were not used in the array experiment, however data from one experiment is shown. Genes upregulated by Merlin expression are shown in red, downregulated in blue.

**Figure S4.**
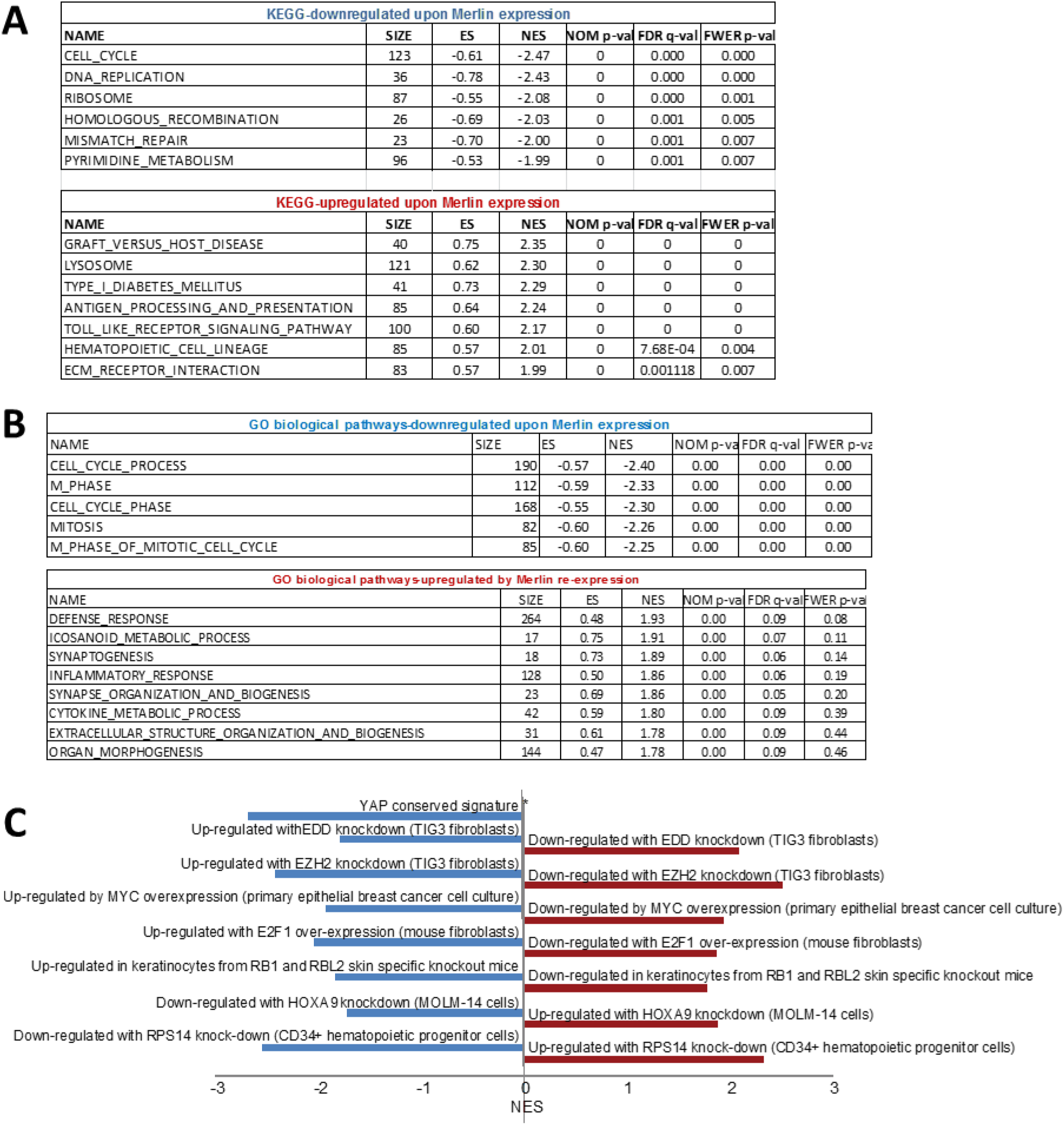
Gene Set Enrichment Analysis of microarray data. (A-B) Gene Set Enrichment Analysis (GSEA) of Merlin re-expression microarray results compared by KEGG (A) or GO biological pathways (B).Top 5-8 up- and down-regulated gene set pathways shown. (C) Selected transcriptional signatures from Oncogenic Signatures database associated with Merlin hypersensitivity by GSEA. * note that YAP conserved signature only contains genes upregulated by YAP expression. NES, normalized enrichment score.

**Figure S5.**
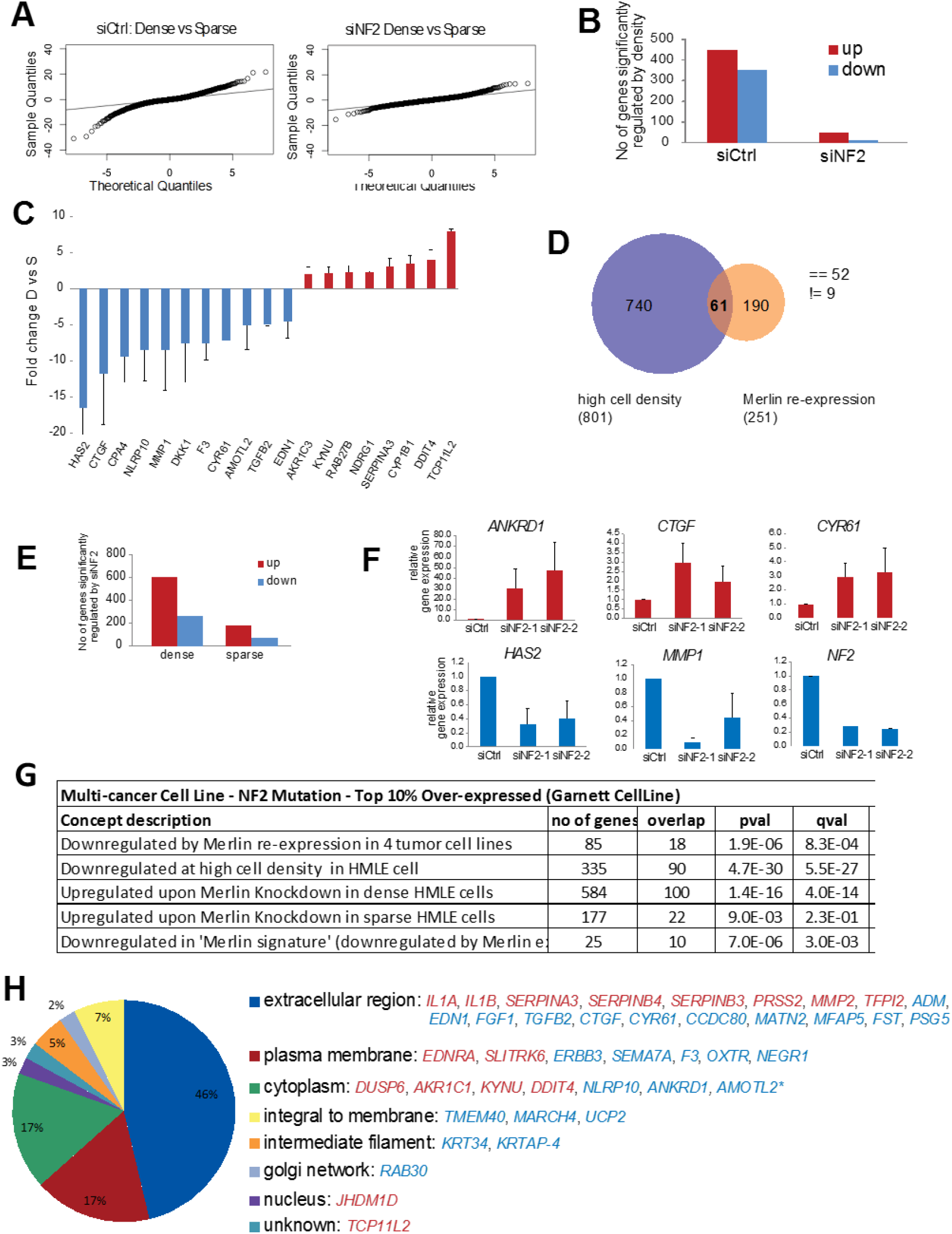
Merlin is involved in the transcriptional response to high cell density. **(A)** Gene expression changes in dense (D) and sparse (S) HMLE cells 3 days after transfection with scramble (Ctrl) or NF2 siRNAs were analyzed by microarray. siControl cells show greater dispersion of limma t-values compared to the theoretical null distribution (black line). **(B)** As in A but showing number of significantly regulated genes (p≤ 0.05, fold-change >1.5). **(C)** Validation by RT-qPCR of representative genes regulated by cell density in HMLE cells. Fold change in dense compared to sparse cells. Error bars indicate SD from n=3. **(D)** Overlap between the Merlin re-expression and high cell density signatures. The majority of genes (85%, 52 of 61) are regulated in the same direction (==), while 9 are regulated in the opposite manner (!=). **(E)** Merlin knockdown has a greater effect on gene expression in dense cells. Number of significantly regulated genes (q≤ 0.05, fold change >1.5) is shown. **(F)** Validation by RT-qPCR of representative genes regulated by Merlin knockdown in dense HMLE cells with two independent siRNAs. Gene expression relative to GAPDH and fold change normalized to siCtrl. Error bars indicate SD from n=3. **(F)** Merlin and high cell density signatures detect *NF2* mutational status in the Garnett *et al* multi-cancer cell line study using the Oncomine database. Significance and overlap shown. **(H)** Merlin core gene signature is enriched in extracellular and plasma membrane proteins. Merlin core signature is defined as genes regulated in the opposite direction upon Merlin re-expression in 4 tumour cell lines and knockdown in dense HMLE cells. Red indicates positively regulated by Merlin (downregulated by knockdown and upregulated by re-expression), blue negatively regulated by Merlin (upregulated by knockdown and downregulated by re-expression). Protein location based on Gene ontology.

**Figure S6.**
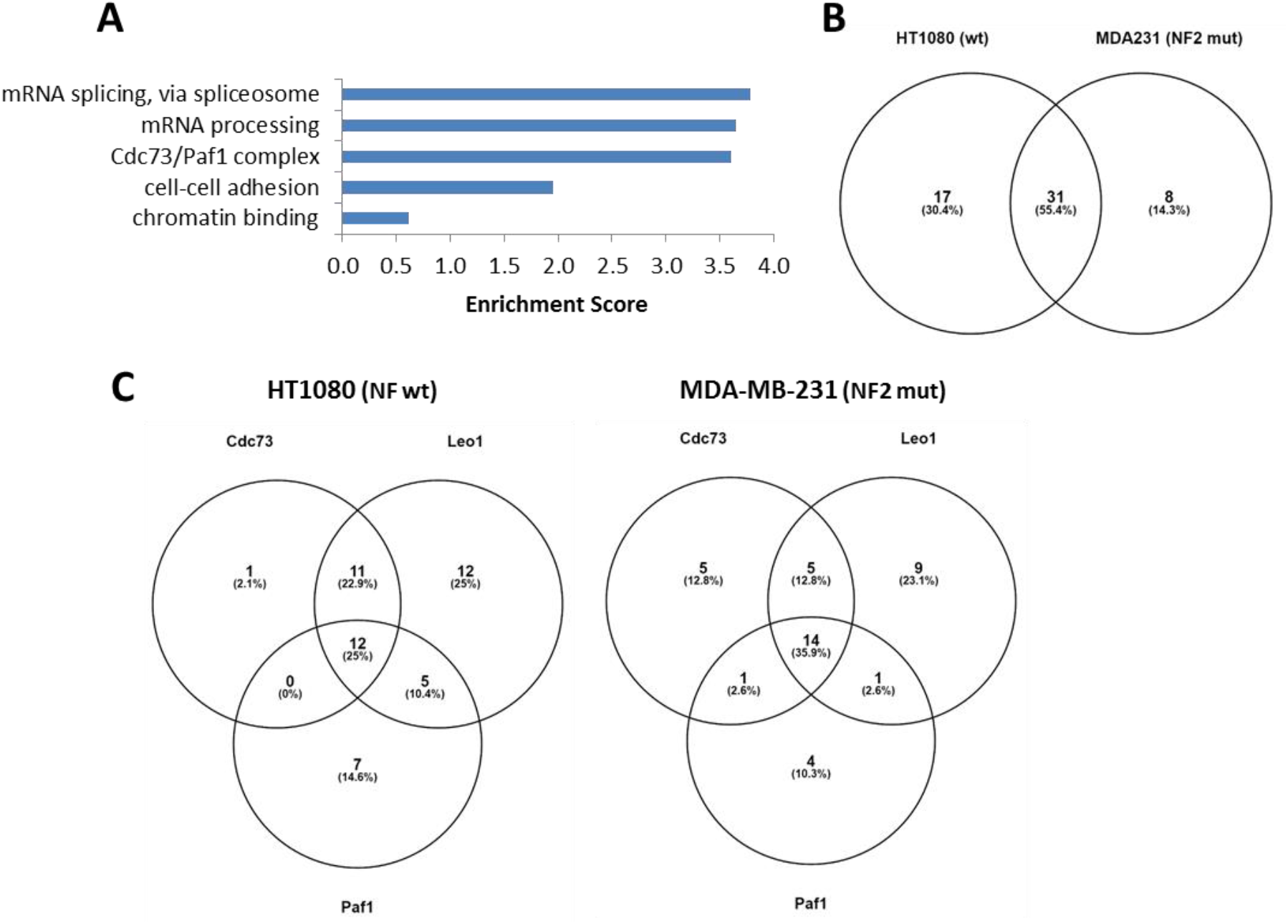
Proteomic analysis of PAF1, CDC73 and LEO1 subunits of PAFC. (A). Gene Ontology term enrichment of proteins co-purifying with PAFC subunits. TAP6-CDC73, PAF1 and LEO1 were expressed and affinity purified from either HT1080 (Merlin wild type) or MDA-MB-231 (Merlin deficient) cells. Co-purifying proteins identified by mass spectrometry were subjected to functional annotation analysis with GO biological process, cellular component and molecular function terms using DAVID (Huang da et al., 2009). Terms correspond to GO:000368, GO:0098609, GO:0016593, GO:0006397 and GO:0000398. (B). Venn diagram showing overlap of proteins identified in HT1080 and MDA-MB-231. See Table S11 for full lists. (C). Venn diagram of proteins identified by AP-MS with each bait (CDC73, LEO1 and PAF1) in either HT1080 or MDA-MB-231 cells.

**Figure S7.**
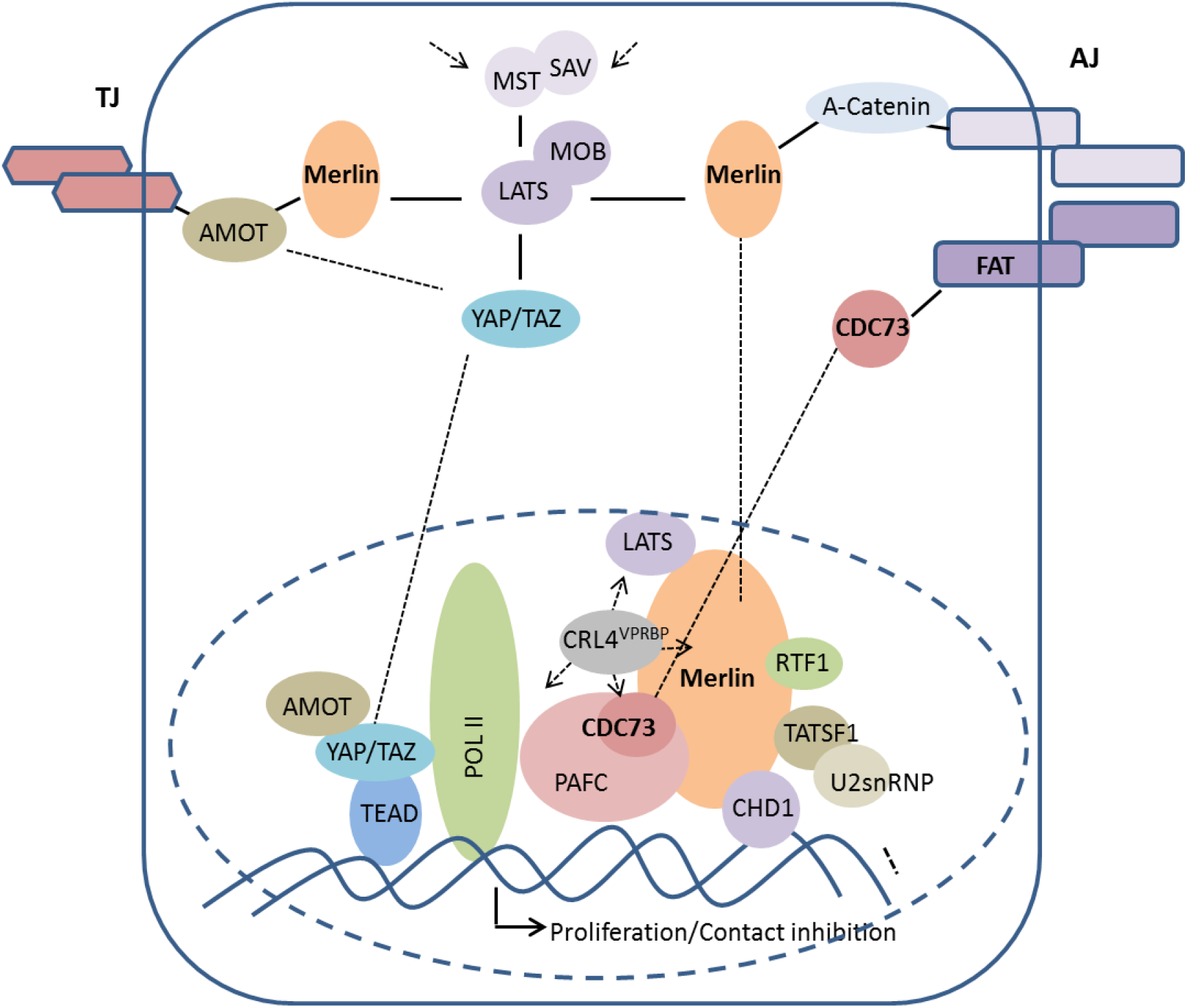
Model of Merlin’s dual role within the Hippo pathway to coordinate different steps in the transcription cycle of genes involved in contact inhibition. At the membrane, Merlin associates with cell-cell contact associated proteins such as α-catenin at adherens junctions and AMOT proteins at tight junctions whereas FAT cadherins may associate with the CDC73 subunit of the PAF complex at distinct adherens junctions. Merlin at the membrane primarily functions to regulate LATS-dependent phosphorylation and inactivation of YAP/TAZ. Upon Merlin inactivation (e.g. upon loss of cell-cell contacts), YAP/TAZ translocates to the nucleus to mediate TEAD dependent recruitment of Pol II and transcription initiation. In the nucleus, Merlin interacts with the CRL4^VPRBP^ E3 ubiquitin ligase (Li et al., 2012) and with proteins involved in transcription elongation and RNA processing including the PAFC, CHD1, RTF1 and TAT-SF1 (this study). Through these interactions, Merlin regulates post-initiation events such as pause release, elongation rate and/or RNA processing in at least a subset of target genes. Merlin could use CRL4^VPRBP^ mediated ubiquitination of some associated proteins (such as CDC73 and/or other CRL4^VPRBP^ substrates) to regulate their properties and/or function within this macromolecular complex. AMOT also has dual membrane/cytoplasm and nuclear roles and is required for YAP transcriptional activity of some target genes (Hong, 2013; Ragni et al., 2017). ZO-2, another tight-junction associated protein also interacts with YAP in the nucleus (Oka et al., 2010). It is therefore possible that separate membrane/cytoplasm and nuclear functions of several of its components is a common feature of the Hippo pathway that allows for independent layers of regulatory control. Note that reported additional interactions of FAT with Hippo pathway components are not shown (Martin et al., 2018; Ragni et al., 2017) but also suggest multiple layers of regulation of the Hippo pathway by FAT proteins. Merlin mays also cooperate/antagonize with transcription factors other than YAP/TAZ (not shown). AJ: adherens junctions, TJ: tight junctions.

## Supplementary Tables (Excel files)

Table S1. Genes regulated by Merlin expression in Merlin wild type cells

Table S2. Genes regulated by Merlin expression in NF2 mutant cells

Table S3. Genes regulated by high cell density in control siRNA HMLE cells

Table S4. Genes regulated by high cell density in siNF2 HMLE cells

Table S5. Genes regulated in the same direction by Merlin re-expression in 4 NF2 mutant cell lines and high cell density in HMLE cells

Table S6. Genes regulated by NF2 knockdown in dense HMLE cells

Table S7. Genes regulated by NF2 knockdown in sparse HMLE cells

Table S8. Merlin core gene signature

Table S9. Genes where Pol II association is regulated by wild type vs L46R Merlin expression in MDA-MB-231 by ChIP-seq

Table S10. Genes where LEO1 association is regulated by wild type vs L46R Merlin expression in MDA-MB-231 by ChIP-seq.

Table S11. Proteins identified by mass spectrometry with TAP6-CDC73, -PAF1 or –LEO1 in HT1080 or MDA-MB-231 cells

Table S12. Proteins identified by mass spectrometry after affinity purification of TAP6-CDC73 in HEK293T cells co-expressing either Ezrin or Merlin

## SUPPLEMENTARY METHODS

### Antibodies and constructs

AMOTL1 (HPA001196), Flag (M2), FAT 1 (HPA001869) and HA-peroxidase antibodies were obtained from Sigma. Merlin (E-2 and A-19), RNA Pol2 (N-20), SPT16 (H-4), HA (Y-11), MYC (A-14) and anti-goat-HRP (sc-2020) antibodies from Santa Cruz and p21 (05-345) from Millipore. AMOT (303A), CDC73 (170A), CHD1 (218A), CTR9 (395A), DDB1 (462A), LEO1 (175A), Merlin iso1 (A578), Merlin iso2 (A579), PAF1 (172A), PELP1 (180A), RTF1 (179A), YAP (309A), SF3A3 (507A), SF3B1 (997A), SF3B3 (508A), TAT-SF1 (023A), VPRBP (887A and 888A), FAT 1 (402A and 403A) antibodies were from Bethyl Laboratories, FLAG-Dylight680 and HA-IRDye800 from rockland. Anti-mouse-HRP (NA 931) and anti-rabbit-HRP (NA934) were obtained from Amersham. cDNAs were cloned into the pENTR vector (Invitrogen) and transferred to Gateway-compatible expression vectors using recombination-mediated Gateway technology (Invitrogen). Mutations were generated using the QuikChange site-directed mutagenesis kit (Stratagene).

### Cell culture and virus generation

Dulbecco’s Modified Eagle’s Medium (DMEM) and Roswell Park Memorial Institute (RPMI) Medium was supplemented with 10% FBS; DMEM F12 was supplemented with 5% HS, 20 ng/ml EGF, 0.5 μg/ml hydrocortisone, 100 ng/ml cholera toxin and 10 μg/ml insulin. All cells were grown in DMEM except for 786-0, 8505C, ACHN, B-CPAP, CAKI-1, CGTH-W-1, S-117 and SN12C which were grown in RPMI, and HMLE and MCF10A which were grown in DMEM F12. Lentiviruses were generated by transient transfection of HEK293T cells with the lentiviral construct, pMD.G (VSV-G) and p8.91 (gag-pol) packaging vectors. Virus-containing medium was harvested 48 h and 72 h after transfection and supplemented with 5 μg/ml Polybrene (hexadimethrine bromide, Sigma-Aldrich). Cells were transduced with lentivirus and selected 24 hours later with 2.5 mg/ml puromycin (Sigma-Aldrich) or 10 mg/ml Blasticidin (Melford). Retroviruses were generated by transient transfection of the Phoenix ecotropic cell line and virus was collected and cells infected as above. Cell lines expressing the ecotropic receptor EcoR where generated by transduction with amphotropic EcoR retroviruses and selection with Blasticidin. Retroviruses and lentiviruses were titered by infection with serial dilutions in HEK293T and/or IOMM-Lee cells.

### Transfections

2×10^6 HEK293T cells were transfected with 2ug plasmid DNA and 1 mg/ml polyethylenimine (PEI, Polysciences) mixed at a 1:5 ratio in OptiMEM (Gibco). After 30 min incubation at RT the transfection solution was added to cells cells seeded 2 h prior. Fresh medium was added 16-24 h after transfection and cells harvested the day after. For RNAi knockdown experiments cells were seeded 2 h prior transfection using Lipofectamine RNAiMAX (Invitrogen) according to the manufacturer’s recommendations using 10 nM siRNA oligos and lysed 3 days later. Stealth RNAi Negative Control Medium GC Duplex (Invitrogen) was used as control oligo. Stealth siRNA oligos were from Invitrogen and are listed below. Smartpools were obtained from Thermo Scientific.

### Purification of recombinant proteins and ‘Fishing’ experiments

Baculoviruses expressing Merlin wild type and variants expressed as GST-fusions proteins were generated using the FastBac method (Invitrogen). 3 days after infection, Sf9 cells were lysed in PBS-E LB and clarified lysates purified with Gluthathione Sepharose beads as above. After extensive washing, beads and bound proteins were stored in 50% glycerol at −20 °C. 10 μl of beads were added to lysates, incubated for 2 h, washed and eluted with LDS sample buffer. Co-purifying proteins were detected by Western blotting with the appropriate antibodies.

### Nuclear fractionation/ enrichment

HT-1080 fibrosarcoma cells were infected with lentiviruses expressing shRNA hairpins targeting non-silencing control, NF2, VPRBP or PELP1 and selected in 2.5 μg/ml puromycin. shRNAmir sequences were from pGIPZ library (Open Biosystems):

Non-silencing: CTCTCGCTTGGGCGAGAGTAAG

NF2-1: CCGGTGTGAACTACTTTGCAAT

NF2-3: CGCAGCAAGCACAATACCATTA

VPRBP: CGCACTTCAGATTATCATCAAT

PELP1: AAGGATTTGACAGTTATTAATA.

Cells were washed twice with cold PBS, washed once quickly with hypotonic buffer without Triton X-100 and lysed in hypotonic buffer (20 mM Hepes (pH 7.5), 10 mM NaCl, 1.5 mM MgCl2, 1 mM EDTA, 0.1% (v/v) Triton X-100, 20% Glycerol (v/v)) on ice for 10 min. Cells were scraped and spun at 150 g for 10 min at 4°C. The supernatant was kept as the cytosolic fraction. The nuclear pellet was washed once in hypotonic buffer without Triton X-100 but supplemented with 1 mM DTT, phosphatase inhibitor (Halt, Thermo Scientific) and protease inhibitor (Roche) and lysed in 5x pellet volume of hypertonic buffer (20 mM Hepes (pH 7.5), 0.5 M NaCl, 1.5 mM MgCl2, 1 mM EDTA, 20% Glycerol (v/v)) for 30 min at 4°C. The lysate was subsequently spun at 16000 g for 10 min. The supernatant yielded the nuclear fraction. The pellet was lysed in 20 μl 2x LDS for nuclear insoluble fraction.

### Gene expression analysis

Gene expression profiles were determined using Whole-transcript Expression Analysis on GeneChip^®^ Human Gene 2.0 ST Arrays (Affymetrix). Total RNA was extracted 48hours after lentiviral infection using the RNeasy Mini Kit (Qiagen). cDNA was generated using the WT Expression Kit (Ambion) and labeled using the GeneChip WT Terminal Labeling Kit (Affymetrix). The hybridisation protocols were performed on GeneChip Fluidics Station 450, and scanned using the Affymetrix GeneChip Scanner. Expression data was pre-processed and normalised using ‘affy’ and the RMA algorithm (Gautier et al., 2004) and differential expression determined using ‘limma’ (Smyth, 2005). Unless otherwise stated, significant difference of expression is taken as q < 0.05 (false discovery rate adjusted) and fold-change greater than +/− 1.5.

For qRT-PCR, RNA was reverse transcribed using Protoscript M-MuLV First Strand cDNA Synthesis Kit (New England BIolabs) and analysed by quantitative real-time PCR (qPCR) with SYBR green fluorescence. Data were normalized to Actin and GAPDH expression as indicated. See supplementary experimental procedures for primer sequences.

Microarray data has been deposited in GEO as GSE53501 and ChIP-seq. data in EMBL-EBI as SRP035339.

### ChIP-qPCR and ChIP sequencing

Cells were infected with lentiviruses expressing Ezrin or Merlin, selected the next day with puromycin for 24 hours and fixed with 1% formaldehyde 48 hours after infection. DNA was sheared by sonication (Vibra-cell, Sonics) to an average length of 400 bp in ChIP lysis buffer (1% SDS, 10 mM EDTA, 50 mM Tris-HCl, pH 8.0). Sheared chromatin was diluted 1/10 in ChIP IP buffer (0.1% SDS, 1.1% Triton X-100, 1.2 mM EDTA, 167 mM NaCl, 16.7 mM Tris-HCl, pH 8.0) and protein/DNA complexes were immunoprecipitated over night with indicated antibodies. After extensive washing and decrosslinking DNA was purified by phenol/chloroform extraction and quantified by real-time PCR (Mastercycler ep Realplex, Eppendorf) measuring an increase in SYBR Green (Fermentas). qPCR conditions were the following: 95 °C for 10 min followed by 40 cycles of 95 °C for 15 sec and 60 °C for 1 min. The amount of DNA recovered was normalized against input DNA. See below for primer sequences.

For ChIP-Sequencing cells were fixed, processed and the chromatin IPed as described above. The ChIP-seq libraries were prepared using 300ng of input DNA and ChIP DNA using the NEBNext ChIP-Seq Library Prep Master Mix Set for Illumina (New England Biolabs). After end repair and addition of A base to 3’ ends, barcoded adaptors (Mulitplex Oligos for Illumina, New England Biolabs) were ligated to DNA fragments. Following an 18 cycle PCR, DNA fragments of 150-300bp were purified by agarose gel purification. ChIP-Seq libraries were then normalised, pooled and sequenced on one lane of the HiSeq 2000 platform (Illumina) using the latest v3 reagents, at a final concentration of 12pM. Illumina’s CASAVA software (v1.8.2) was then used to demultiplex the pooled sequence data and create fastq files for each sample.

Raw paired end Illumina data (Illumina pipeline 1.3+) was first trimmed to 50bp then aligned to the Hg19 human reference using BWA version 0.5.9 (Li and Durbin, 2010). Read counts were then generated for each gene in a sample by selecting the longest known transcript and an additional 400bp upstream of the TSS. Further processing was then undertaken using the R statistical programming language (version 2.14.2) (R Development Core Team, 2012) including the DESeq bioconductor package (version 2.11) (Anders and Huber, 2010).

After normalising by library size the antibody control count from each gene was deducted from each sample. Differential enrichment statistics were then calculated using the DESeq package allowing for pooling of samples to compensate for lack of replicates. RNA Pol2 and LEO1 samples were compared independently. All plots were created using the NGSplot graphing toolkit (Liu X., 2012). Data were visualized at representative loci using UCSC genome browser. Data has been deposited in EMBL-EBI as SRP035339.

### Statistical analysis

Data are represented as mean values and the error bars indicate ±SD. Groups were compared by two-tailed unpaired t-test. The significance is indicated as *** for P<0.001, ** for P<0.01 and * for P<0.05.. For expression data multiple testing correction (q-values) were calculated using the Benjamini-Hochberg false dicovery rate (FDR).

### Affinity purification and Mass spectrometry

Tandem affinity purification (TAP) for Merlin was performed from 10 x 15 cm dishes of HEK293T cells, transiently transfected with pcDNA3-TAP6 constructs. For CDC73, 20 x 15 cm dishes of HEK293T stably expressing TAP6-CDC73 (pLEX-MCS, Open Biosystems) and either Merlin or Ezrin (pLentiCMVBlast, addgene 17486) were used. The TAP6-tag contains 3x streptactin, 2x histidine, SBP and FLAG tags. Cells were lysed in PBS-E lysis buffer (LB) Clarified pooled lysates were incubated with Streptavidine Sepharose beads (GE Healthcare) and washed extensively with PBS containing 0.1% TX-100. Bound proteins were first eluted with 2 mM Biotin and then LDS sample buffer. Biotin eluates were TCA precipitated, separated on 4-12% NuPAGE gels (Life Technologies) and stained with SimplyBlue SafeStain (Life Technologies). Gels were cut along the lanes in several pieces, and proteins were digested in-gel with trypsin as described previously (Rosenfeld et al., 1992) (donatello.ucsf.edu/ingel.html). The extracted digests were vacuum-evaporated and resuspended in 10 μl of 0.1% formic acid in water, except the slice containing the bait, that was resuspended in 30 μl. The digests were separated by nano-flow liquid chromatography using a 75 μm x 150 mm reverse phase C18 PepMap column (Dionex-LC-Packings, San Francisco, CA) at a flow rate of 300 nl / min in a NanoLC-1D Proteomics high-performance liquid chromatography system (Eksigent Technologies, Livermore, CA, USA) equipped with a FAMOS autosampler (Dionex-LC-Packings, San Francisco, CA). Mobile phase A was 0.1% formic acid in water and mobile phase B was 0.1% formic acid in acetonitrile. Following equilibration of the column in 5% solvent B, 5 μl of the digests were injected, then the organic content of the mobile phase was increased linearly to 40% over 60 min, and then to 50% in 1 min. The liquid chromatography eluate was coupled to a microionspray source attached to a QSTAR Pulsar mass spectrometer (Applied Biosystems/MDS Sciex, South San Francisco, CA, USA). Peptides were analyzed in positive ion mode. MS spectra were acquired for 1 s in a m/z range between 310 and 1400. MS acquisitions were followed by 3 s collision-induced dissociation (CID) experiments in information-dependent acquisition mode. For each MS spectrum, the most intense multiple charged peaks over a threshold of 30 counts were selected for generation of CID mass spectra. The most common trypsin autolysis products (m/z=523.280+^2^, 421.750+^2^, 737.700+^3^) were excluded. The CID collision energy was automatically set according to mass to charge (m/z) ratio and charge state of the precursor ion. A dynamic exclusion window was applied which prevented the same m/z from being selected for 1 min after its acquisition. Typical performance characteristics were 10000 resolution with 30 ppm mass measurement accuracy in both MS and CID spectra.

A second 5 μl aliquot of the samples was injected using the same chromatographic setting into a hybrid linear ion trap-Fourier transform mass spectrometer (LTQ-FT, Thermo Scientific, San Jose, CA) equipped with a nanoelectrospray ion source. Spraying was from an uncoated 15 μm-inner diameter spraying needle (New Objective, Woburn, MA). Peptides were analyzed in positive ion mode and in information-dependent acquisition mode to automatically switch between MS and MS/MS acquisition. MS spectra were acquired in profile mode using the ICR analyzer in the m/z range between 310 and 1600. For each MS spectrum, the 5 most intense multiple charged ions over a threshold of 200 counts were selected to perform CID experiments. Product ions were analyzed on the linear ion trap in profile mode. CID collision energy was automatically set to 35%. A dynamic exclusion window of 1 Da was applied that prevented the same m/z from being selected for 60 s after its acquisition.

Peak lists from files acquired by both instruments were generated using Mascot Distiller version 2.1.0.0 (Matrix Science, Boston, MA). Parameters for MS processing were set as follow: peak half-width, 0.02; data points per Da, 100. Parameters for MS/MS data were set as follows: peak half-width, 0.02; data points per Da, 100. The peak list was searched against the human subset of the UniProtKB database as of December 16, 2008 (containing 167288 entries) using in-house ProteinProspector version 5.2.2 (a public version is available online). A minimal ProteinProspector protein score of 15, a peptide score of 15, a maximum expectation value of 0.1 and a minimal discriminant score threshold of 0.0 were used for initial identification criteria. Carbamidomethylation and acrylamide modification of cysteine; acetylation of the N-terminus of the protein and oxidation of methionine were allowed as variable modifications. Peptide tolerance in searches of QStar data was 100 ppm for precursors and 0.2 Da for product ions; for LTQ-FT data was 30 ppm for precursors and 0.6 Da for product ions, respectively. Peptides containing two miscleavages were allowed. The number of modification was limited to two per peptide. Carbamidomethylation and formation of acrylamide adducts of cysteine, N-acetylation of the N-terminus of the protein, oxidation of methionine, phosphorylation of serine, threonine or tyrosine, and acetylation of lysine were allowed as variable modifications.

Hits were considered significant when three or more peptide sequences matched a protein entry and the Prospector score was above the significant level. For identifications based on one or two peptide sequence with high scores, the MS/MS spectrum was reinterpreted manually by matching all the observed fragment ions to a theoretical fragmentation obtained using MS Product (Protein Prospector) (Clauser et al., 1999) Number of unique peptides reported combines the results of the searches for both sets of data. Normalized spectral counts were calculated normalizing the total number of spectra acquired for each protein (combining data from both instruments) and normalizing by the length of each polypeptide.

### CDC73 interactome

The final interactome list contained proteins that were identified with at least 2 unique peptides with the CDC73 bait but not an unrelated bait purified in parallel. Label-free quantification was estimated by spectral counts that were normalised to the CDC73 bait. A normalisation factor of 15% was calculated (Merlin-CDC73/Ezrin-CDC73=1.15) and used to adjust the spectral counts of the Ez pulldown. A Merlin/Ezrin fold change was calculated for the proteins detected in both pulldowns. Proteins detected in one pulldown only with at least 3 peptide counts were defined as upregulated (absent in Ez) or downregulated (absent in NF2) to a higher degree than if detected in both pulldowns. The protein list was loaded into DAVID (Huang da et al., 2009) for functional annotation analysis with Gene Ontology (GO) terms. Top ranked enriched functional clusters of GO terms were selected and used to fully annotate the protein interactome list within the QuickGO browser for Gene Ontology terms and annotations. The protein list was then loaded into the STRING database of functional protein association networks where textmining, experiments, and databases were used as active interaction sources to generate a network at medium confidence. The network was exported and loaded into Cytoscape 3.3.0 for integrating the network with attribute data (fold change regulation and functional terms) and visualizing these.

### Oligos

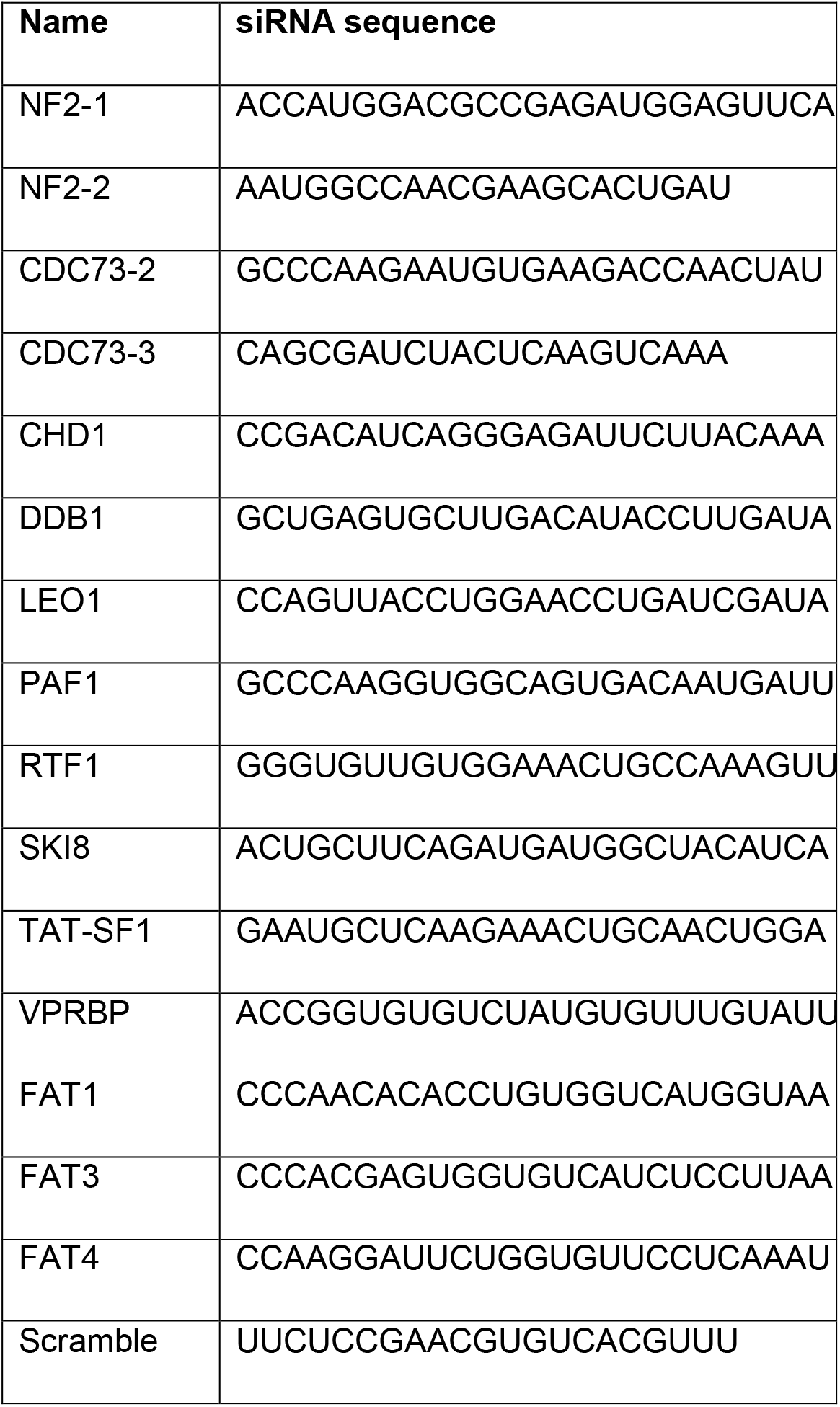

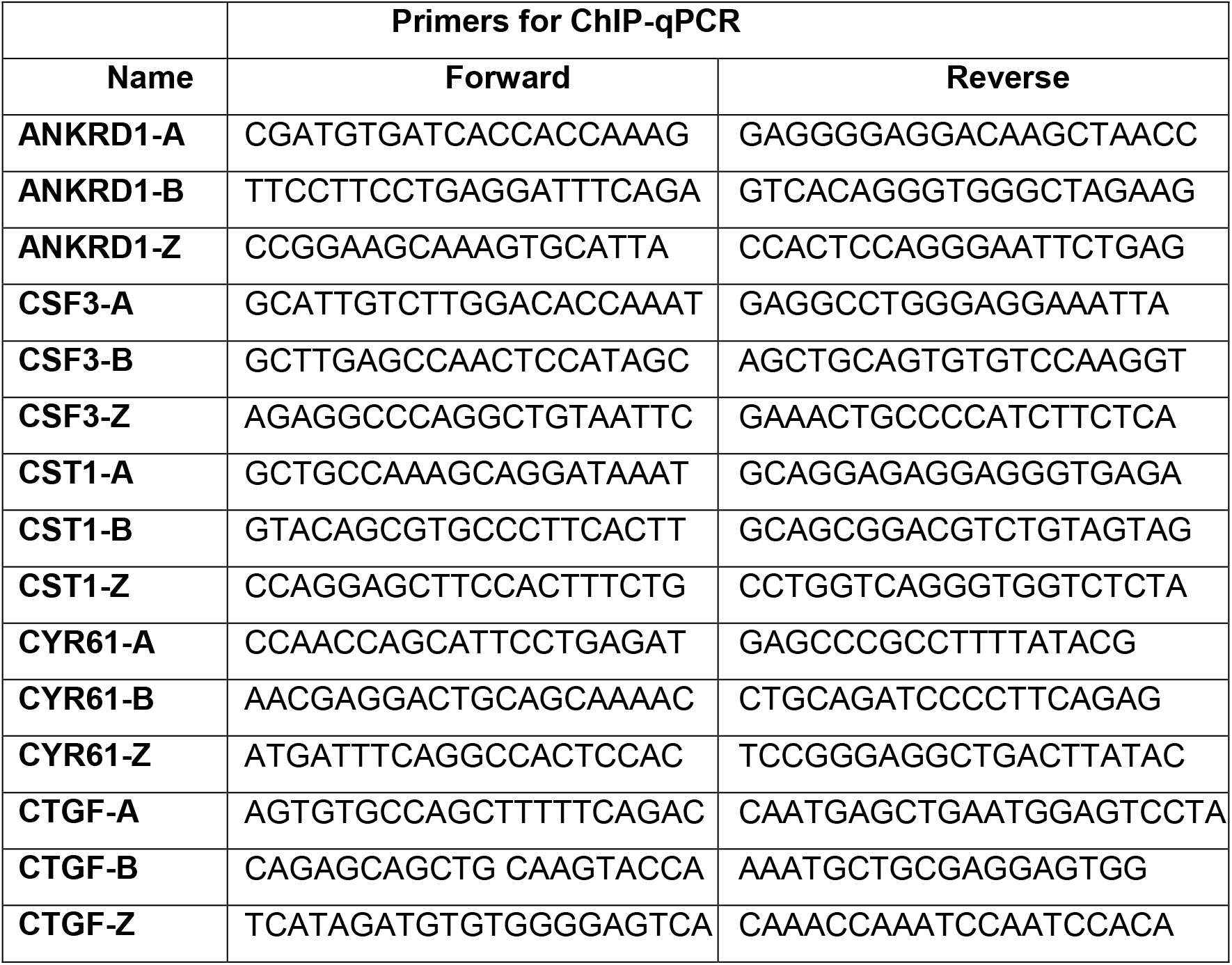

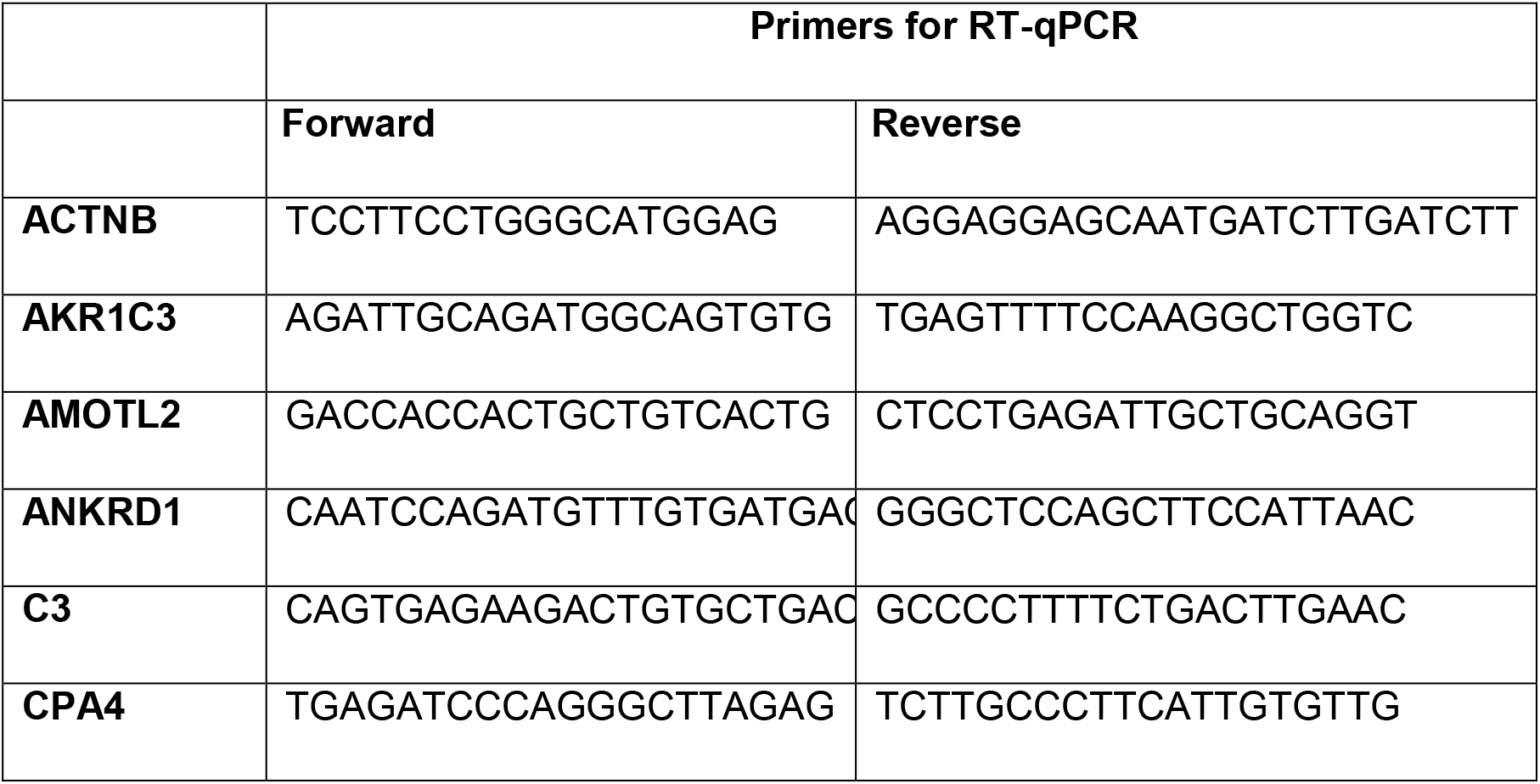
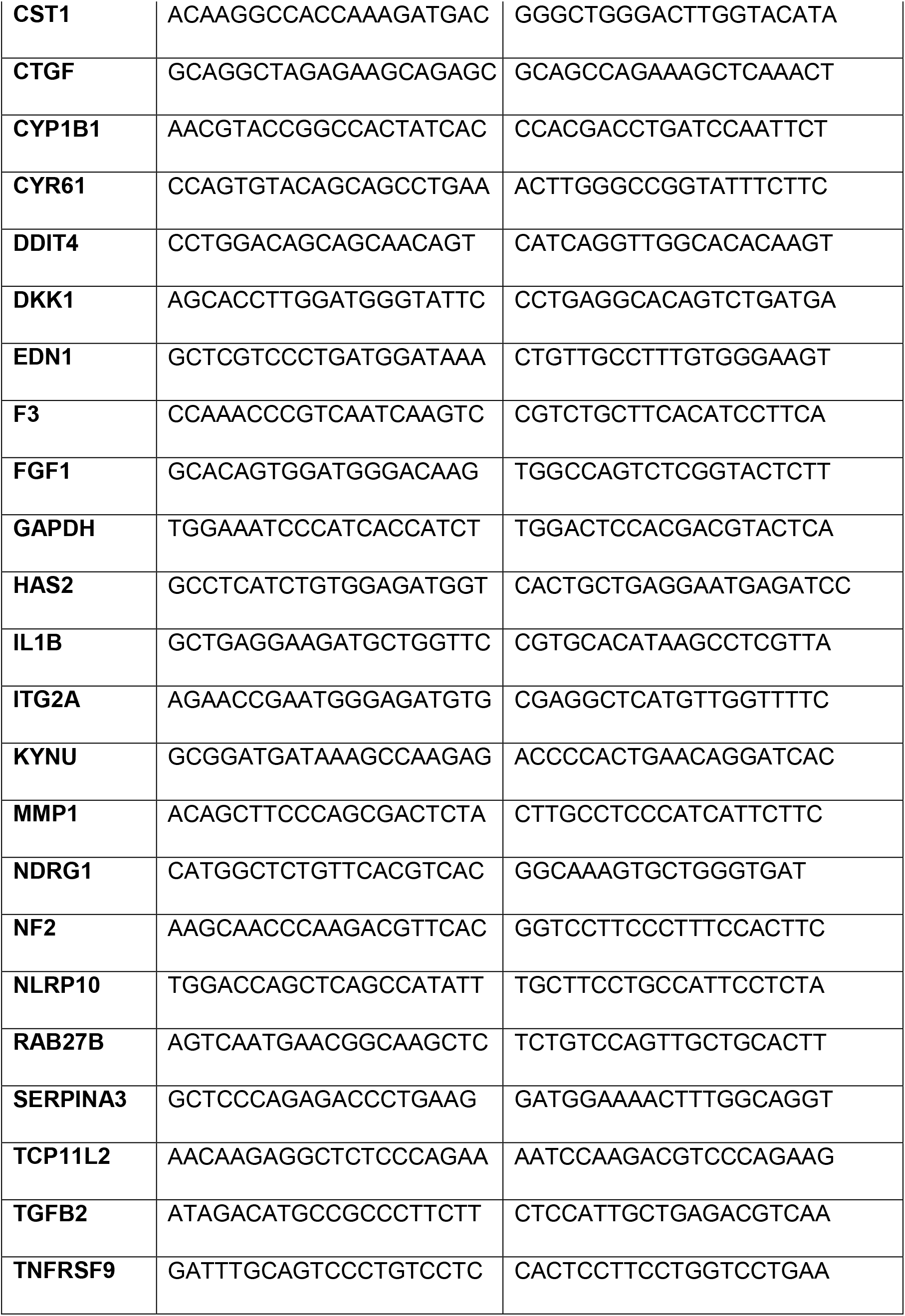

